# Causal evidence for a shared mechanism linking language and tool use via the putamen

**DOI:** 10.64898/2026.02.06.704290

**Authors:** Zhiyu Fan, Haojie Wen, Zaizhu Han, Xiaosha Wang, Yanchao Bi

## Abstract

Both human language and tool use—two hallmark capacities of human cognition—depend on organizing discrete elements, i.e., symbols and actions, into highly constrained structured sequences to achieve a functional goal. However, the neural mechanism linking these capacities is unclear. We combined brain lesion analysis, developmental contrast, and functional neuroimaging to test whether the basal ganglia play a causal role in their shared capacity. In 100 adults with focal brain injury, damage to the putamen disrupted both sentence processing and tool use, with impairments specifically explained by reduced goal-dependent sequence integrity for both tasks. Further comparing populations with typical and deprived early language experience (congenitally deaf adults with vs. without early sign language exposure), we found that early language acquisition was associated with improved tool-use performance and strengthened putaminal responses to such goal dependency, which mediated the relationship between sentence sequence integrity and tool behavior. Together, these results identify the putamen as a key neural substrate supporting goal-dependent sequence integrity across language and action, and show how early language experience shapes this conserved control system.

**One-Sentence Summary:** Language and tool use share a putamen-based mechanism that supports goal-dependent sequence integrity, and this mechanism is strengthened by early language experience.

## Introduction

Language and tool use are defining features of human intelligence. Although they differ in form—language is symbolic composition, tool use is physical manipulation—both require the ability to assemble elements into highly structured sequences. For language, syntax combines words into structured sequences with dependencies spanning varying distances (Gibson, 1998; Hauser et al., 2002; Jackendoff, 2003); skilled tool use integrates multiple objects (hand, tool, target object) and sequential motor acts (e.g., reaching, grasping, and applying force) into goal-directed behaviors constrained by the tool’s learned function and physical affordances, sometimes referred to as “tool syntax” (Martins et al., 2019; Miller et al., 2018; Pastra & Aloimonos, 2012; Roy & Arbib, 2005; Steele et al., 2012). This parallel has long led to the postulation that language and tool use may share a deep computational architecture or even an evolutionary origin (Arbib, 2011; Iriki & Taoka, 2012; Stout & Chaminade, 2012; Vaesen, 2012).

Behavioral and neural evidence has been reported to support this notion. Tool proficiency—not manual dexterity—predicts syntactic fluency (Brozzoli et al., 2019), and training in one domain transfers bidirectionally to the other (Py et al., 2025; Thibault et al., 2021). Relatedly, language instruction improves the fidelity of stone-tool knapping skill transmission (Cataldo et al., 2018; Lombao et al., 2017; Morgan et al., 2015). At the neural level, language and tool processing activate partially overlapping frontoparietal and basal ganglia regions (Fazio et al., 2009; Goldenberg & Randerath, 2015; Higuchi et al., 2009; Vingerhoets et al., 2013; Weiss et al., 2016). A recent study further demonstrated cross-domain neural representational similarity in the basal ganglia during tool-use planning and syntactic processing in sentence comprehension (Thibault et al., 2021), suggesting that both domains may recruit overlapping neural representations. Within language and tool processing, various roles of these regions have been discussed, including semantics, phonology and syntax for language (Bocanegra et al., 2015; Copland et al., 2021; Friederici, 2003; Friederici & Kotz, 2003; Moro et al., 2001; Tan et al., 2026; Tzourio-Mazoyer et al., 2002; Ullman, 2001), and semantic knowledge, action imitation, action planning and organization for tools (Caspers et al., 2010; Choi et al., 2001; Goldenberg & Spatt, 2009; Graybiel, 1998; Johnson-Frey, 2004; Martin, 2007). The overlap invites further articulation and causal tests of the neural operations shared by these capacities, more so than with other cognitive processes, and such causal tests are currently missing.

Here we examine one possible intersection—the extent to which the attainment of a higher-order goal imposes constraints on a specific sequence. While many behaviors unfold sequentially, goal dependencies require choices to be selected and sequenced with respect to outcomes that are not locally available at the time of action or interpretation. For instance, it has been shown that compared with general action events, where the order of some elements is flexible, the constraints on such sequencing in sentence processing are particularly rich (Coopmans et al., 2023). In language, the broader goal of conveying an intended message with a selected syntactic structure constrains the word sequences (e.g., Bock & Levelt, 1994; Gibson, 1998; Giglio et al., 2024). This property of language sequence is also applicable to tool use. In tool use, higher-order functional goals constrain early grasp configurations and intermediate sub-actions, such that initial motor choices are selected with respect to the biomechanical and functional requirements of the intended outcome, as opposed to local optima. A classical contrast is that grasping a pair of scissors to cut paper (functional grasp) is different from grasping it to move (Buxbaum et al., 2003; Garcea & Buxbaum, 2019; Rosenbaum et al., 1990, 2001; see Fig. 3A). Although the representational content differs across domains—linguistic dependencies are abstract and symbolic, whereas tool-use dependencies are grounded in biomechanical and body–effector–object constraints—both may rely on this operation that integrates higher-order goal requirements with sequential choices over time. Quantifying the strength of such goal dependency provides a domain-general measure of sequential properties across symbolic and motor behaviors (Coopmans et al., 2023; Greenfield, 1991; Greenfield & Westerman, 1978).

Brain regions implicated in both language and tool processing—most notably the basal ganglia—are well positioned to support such goal-dependent sequential properties. Beyond language and tool processing, a large body of work has shown that the basal ganglia play a central role in the control of sequential behavior, including selecting among competing alternatives, gating the initiation or suppression of actions, biasing cortical processing based on task goals and expected outcomes, and learning which selections lead to successful outcomes through reinforcement signals (Alexander et al., 1986; Badre & Frank, 2012; Bamford et al., 2018; Chatham & Badre, 2015; Frank et al., 2001; Frank & Badre, 2012; Graybiel, 1998; Jin et al., 2014; Jin & Costa, 2010, 2015; O’Reilly & Frank, 2006; Soni & Frank, 2025; Yang et al., 2025). Through recurrent interactions with cortical regions that encode domain-specific representational content, cortico–basal ganglia circuits may therefore contribute to regulating goal-directed sequencing, which is particularly heavily constrained in sentence and tool actions.

To test these ideas, we combined causal, developmental, and functional approaches (Fig. 1). In Study 1, we used voxel-based lesion–symptom mapping in adults with focal brain injury to identify regions that are jointly necessary for sentence processing and tool use. We examined whether their lesion–behavioral convergence was specifically explained by impairments in processing sequence structures across both domains, after controlling for other domain-specific processes (lexical semantics, action imitation) and other potentially shared components (working memory capacity and broader action semantics). Study 2 leveraged early language deprivation as a developmental causality study within a congenitally deaf population, contrasting individuals with and without early access to natural language (Mayberry et al., 2002, 2011; Newport, 1990; Wang et al., 2023), to test whether early linguistic experience calibrates the neural encoding of goal-dependent sequencing during tool processing.

**Figure 1.**
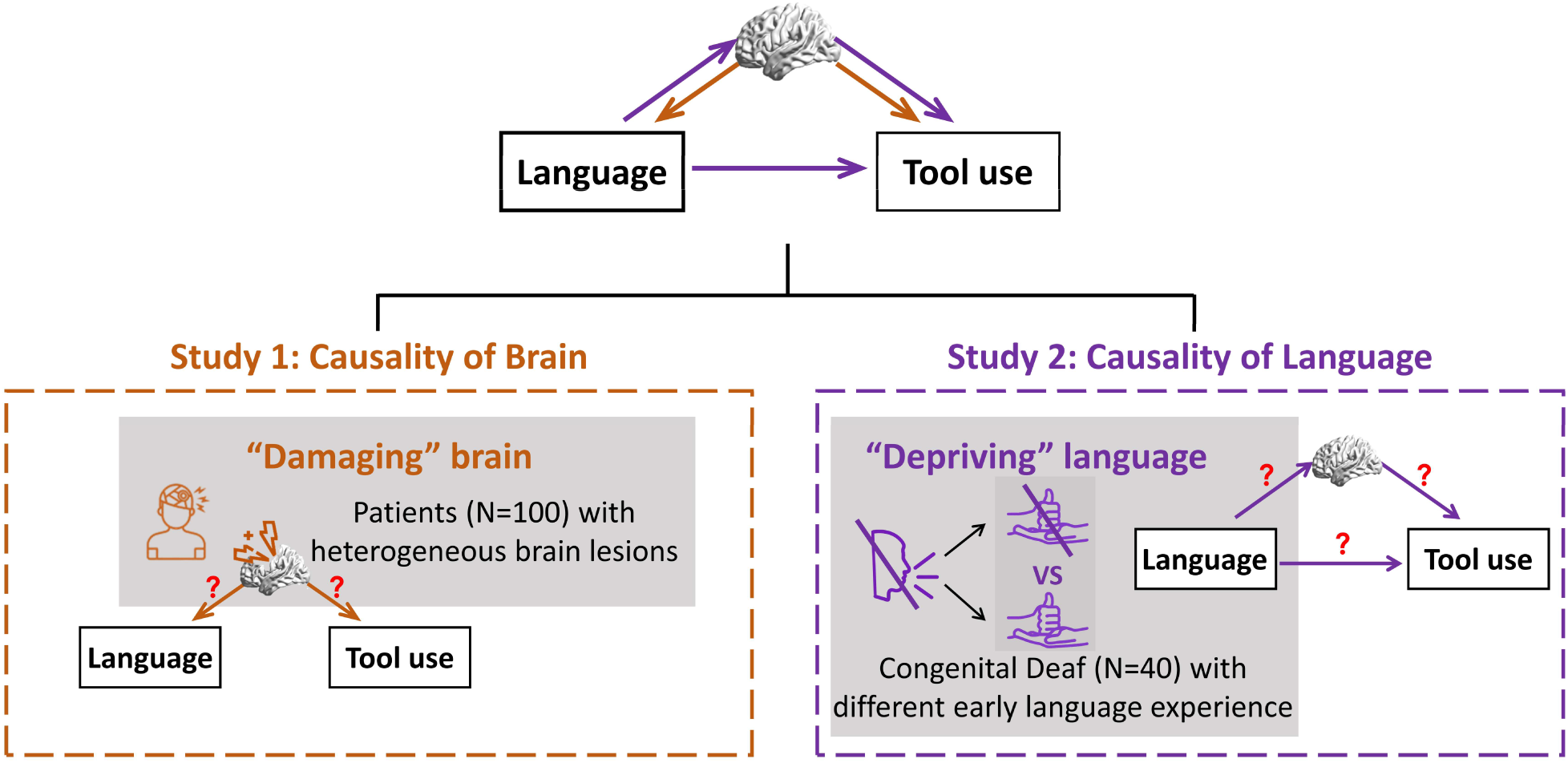
Flowchart of the current study. The schematic illustrates how the two studies jointly investigate the neural causal mechanism—goal-dependency—that supports structured sequence processing in both language and tool use. Study 1 uses a lesion approach, asking whether focal damage to this substrate produces parallel disruptions in language processing and tool-use behavior. Study 2 uses a developmental contrast, and examines whether early language deprivation shapes this shared neural encoding of goal dependency that supports tool use. Together, these complementary approaches provide converging evidence for a shared, causally relevant neural mechanism underlying language and tool cognition.

Together, these studies move beyond correlational overlap to reveal a causal, domain-general neural computation for goal dependency processing in language and tool use, and show how a conserved sequencing circuit across symbolic and motor domains—tuned through early language acquisition—supports the uniquely human capacity to build goal-directed structure from serial experience.

## Results

We implemented a stepwise analytical strategy across two studies to examine goal dependency in language and tool use. In Study 1, we first isolated the structured sequence component of both domains by controlling for other processes relevant to the tasks. Using these residualized behavioral indices, whole-brain voxel-based lesion–symptom mapping (VLSM) revealed overlapping lesion–deficit associations that localized a shared neural substrate. Within this substrate, we then quantified the goal-dependent sequence integrity in each domain. In language, it was indexed by sensitivity to goal-dependent syntactic constraints as reflected in the proportion of syntactically well-formed sentences, whereas in tool use it was indexed by goal-dependent grasp organization. We then tested whether these deficits exhibited voxel-wise pattern alignment within the shared substrate. In Study 2, we probed the neural mechanisms supporting goal-dependency by parametrically modeling the object-specific goal-dependency strength in brain activities elicited while viewing pictures of objects. This allowed us to assess how early language experience shapes tool behavioral performance, neural encoding of goal-dependency, and whether the latter mediates the effect of early language experience on tool-use behavior.

### Causal necessity of the shared neural substrate of the language and tool-use sequence

We first tested whether language and tool use rely on shared neural substrates by performing VLSM in 100 brain-injured patients (Fig. 1, left panel; Fig. 2A, lesion overlay map). Language performance was indexed by sentence comprehension accuracy on a standardized picture–sentence matching task, and tool use was indexed by expert-rated correctness of functional demonstrations across a set of common household tools. Patients were impaired in both sentence comprehension (*t*_(141)_ = –5.74, *p* < 0.001, *d* = –0.86) and tool use (*t*_(141)_ = –11.43, *p* < 0.001, *d* = – 1.68), compared to 43 demographically matched healthy controls. Within the patient group, performance across the two domains was strongly correlated (*r* = 0.53, *p* < 0.001; Fig. S1A), suggesting the shared vulnerability of linguistic and technical abilities.

**Figure 2.**
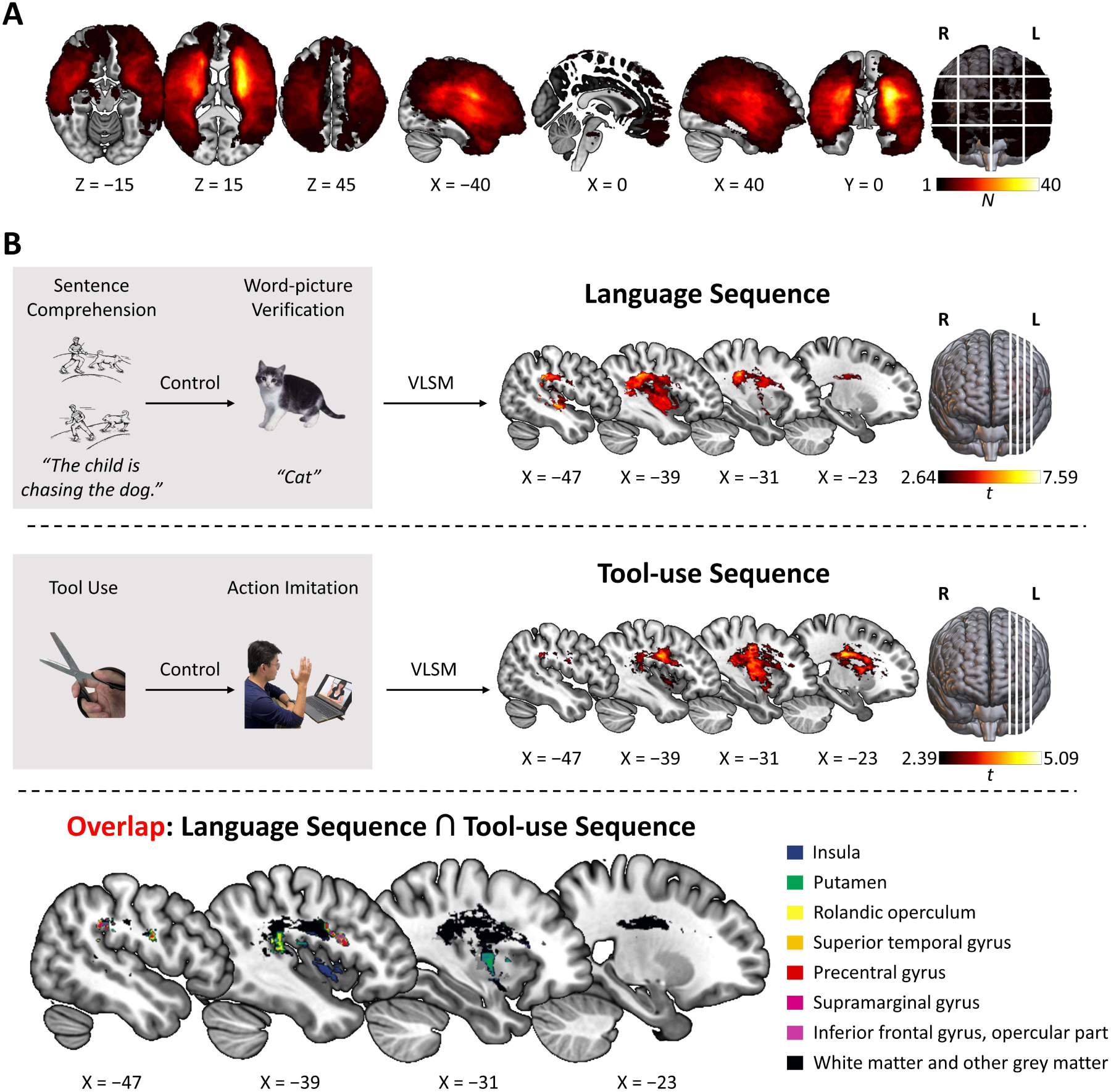
Study 1: The shared neural substrate underlying language sequence and tool-use sequence. ***A***, Lesion overlay map for all 100 patients. The colors indicate the number of individuals showing damage at each voxel. ***B***, VLSM results for language sequence (top panel) and tool-use sequence (middle panel) and their overlap (bottom panel). Residualized indices of language and tool-use sequence were derived from sentence comprehension and tool-use performance by regressing out word–picture verification and action imitation, respectively (voxel-wise FDR corrected, *q* < 0.05). The color bars show voxel-wise t-values. Spatial overlap of the two maps with seven major grey-matter ROIs (≥200 voxels) shown in color. The remaining four grey-matter ROIs (<200 voxels), together with white-matter regions, are shown in black. L denotes the left hemisphere, and R denotes the right hemisphere in all panels.

To isolate sequential components from domain-specific demands in language and tool use, we applied domain-tailored controls in each domain. Given that sentence comprehension integrates both lexical semantics and sentence-level structure building (word combinations), we isolated the structured sequence component by regressing out word-level semantic performance measured with a visual word–picture verification task using non-tool categories. Because tool manipulation inherently involves motor execution, we explicitly separated functional-goal-directed structure formation from peripheral motor ability measured with an imitation task involving non-tool actions. The resulting residualized indices capture the language and tool-use sequences (Fig. 2B), reflecting the combination of discrete elements into a structured sequence. Even after lexical-semantic and motor contributions are removed, the language and tool-use sequences remained correlated (partial *r* = 0.32, *p* < 0.001; Fig. S1B), indicating a higher-order behavioral link grounded in sequential rather than domain-specific, non-sequential semantic or motoric processes.

Whole-brain VLSM revealed overlapping lesion–deficit associations for indices of the language sequence and tool-use sequence in 11 regions of the left hemisphere, including the putamen and precentral gyrus, which were subsequently examined as independent regions of interest (ROIs) in further analyses (FDR *q* < 0.05; Fig. 2B; Table 1; see also Fig. S1C for similar patterns in the raw measures of sentence comprehension and tool use).

**Table 1.**
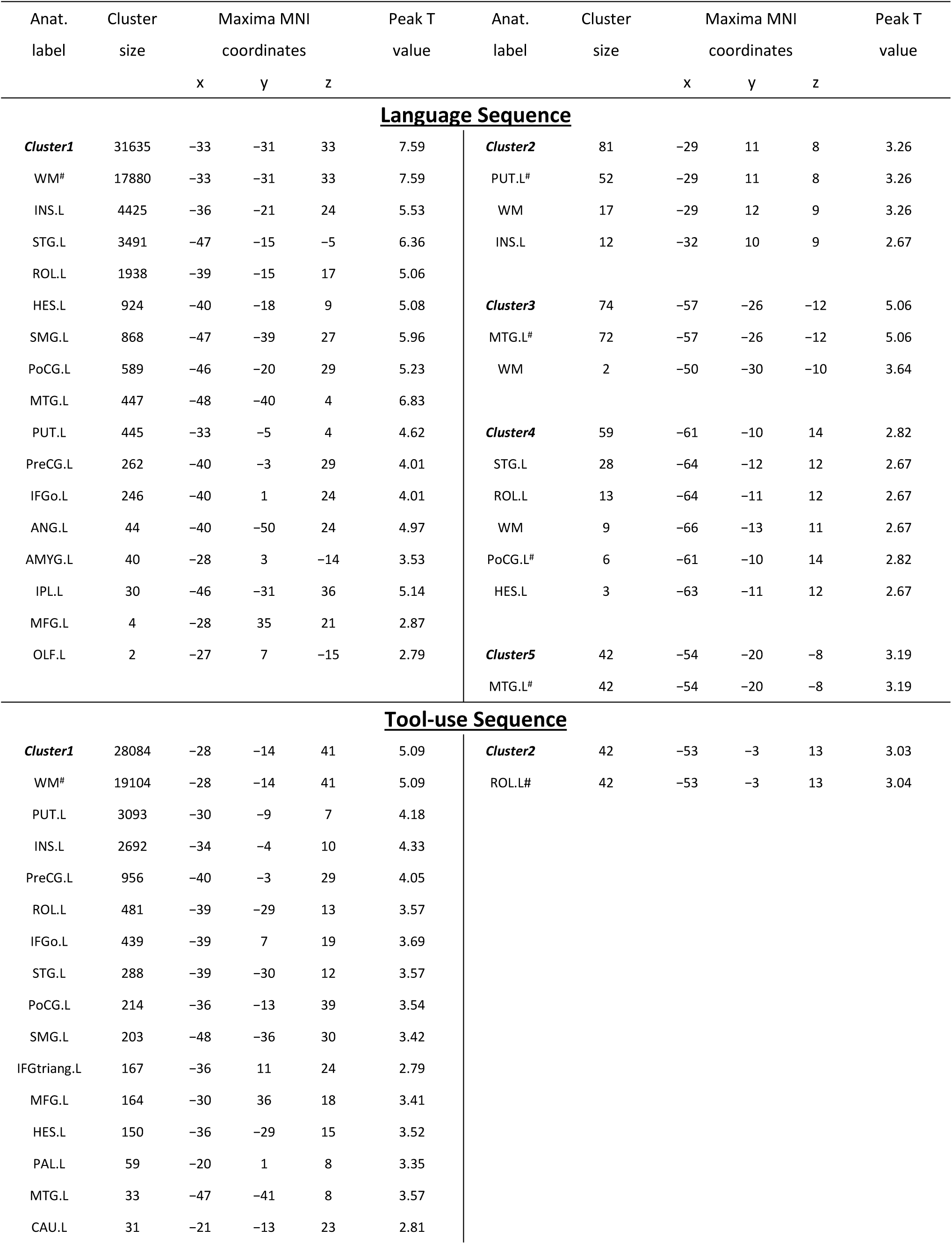
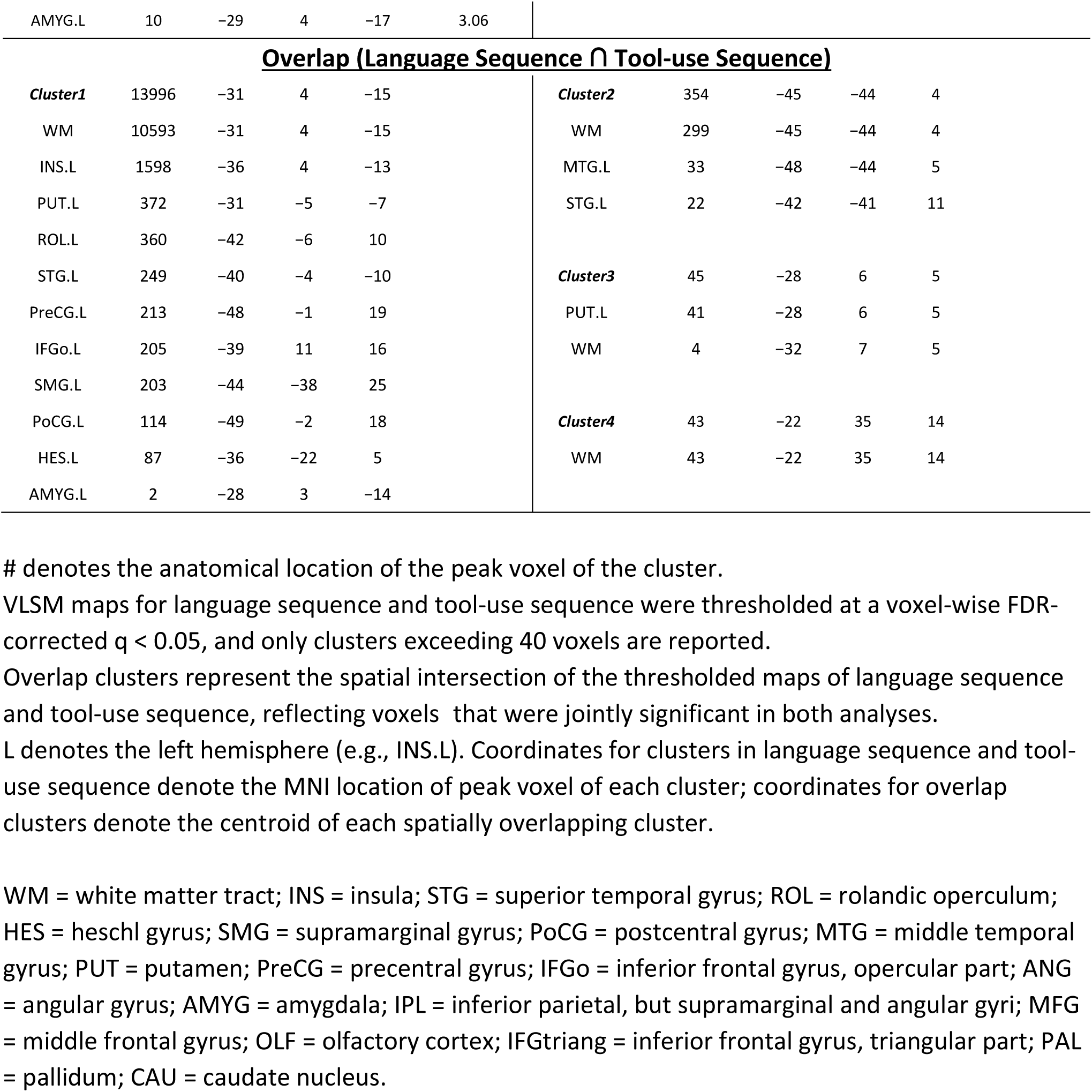
Cluster report of the whole-brain VLSM results for the language sequence, tool-use sequence and their overlap.

| Anat.<br>label | Cluster<br>size | Maxima MNI<br>coordinates |  |  | Peak T<br>value | Anat.<br>label | Cluster<br>size | Maxima MNI<br>coordinates |  |  | Peak T<br>value |
| --- | --- | --- | --- | --- | --- | --- | --- | --- | --- | --- | --- |
|  |  | x | y | z |  |  |  | x | y | z |  |
| **Language Sequence** |  |  |  |  |  |  |  |  |  |  |  |
| **Cluster1** | 31635 | -33 | -31 | 33 | 7.59 | **Cluster2** | 81 | -29 | 11 | 8 | 3.26 |
| WM# | 17880 | -33 | -31 | 33 | 7.59 | PUT.L# | 52 | -29 | 11 | 8 | 3.26 |
| INS.L | 4425 | -36 | -21 | 24 | 5.53 | WM | 17 | -29 | 12 | 9 | 3.26 |
| STG.L | 3491 | -47 | -15 | -5 | 6.36 | INS.L | 12 | -32 | 10 | 9 | 2.67 |
| ROL.L | 1938 | -39 | -15 | 17 | 5.06 | **Cluster3** | 74 | -57 | -26 | -12 | 5.06 |
| HES.L | 924 | -40 | -18 | 9 | 5.08 | MTG.L# | 72 | -57 | -26 | -12 | 5.06 |
| SMG.L | 868 | -47 | -39 | 27 | 5.96 | WM | 2 | -50 | -30 | -10 | 3.64 |
| PoCG.L | 589 | -46 | -20 | 29 | 5.23 | **Cluster4** | 59 | -61 | -10 | 14 | 2.82 |
| MTG.L | 447 | -48 | -40 | 4 | 6.83 | STG.L | 28 | -64 | -12 | 12 | 2.67 |
| PUT.L | 445 | -33 | -5 | 4 | 4.62 | ROL.L | 13 | -64 | -11 | 12 | 2.67 |
| PreCG.L | 262 | -40 | -3 | 29 | 4.01 | WM | 9 | -66 | -13 | 11 | 2.67 |
| IFGo.L | 246 | -40 | 1 | 24 | 4.01 | PoCG.L# | 6 | -61 | -10 | 14 | 2.82 |
| ANG.L | 44 | -40 | -50 | 24 | 4.97 | HES.L | 3 | -63 | -11 | 12 | 2.67 |
| AMYG.L | 40 | -28 | 3 | -14 | 3.53 | **Cluster5** | 42 | -54 | -20 | -8 | 3.19 |
| IPL.L | 30 | -46 | -31 | 36 | 5.14 | MTG.L# | 42 | -54 | -20 | -8 | 3.19 |
| MFG.L | 4 | -28 | 35 | 21 | 2.87 |  |  |  |  |  |  |
| OLF.L | 2 | -27 | 7 | -15 | 2.79 |  |  |  |  |  |  |
| **Tool-use Sequence** |  |  |  |  |  |  |  |  |  |  |  |
| **Cluster1** | 28084 | -28 | -14 | 41 | 5.09 | **Cluster2** | 42 | -53 | -3 | 13 | 3.03 |
| WM# | 19104 | -28 | -14 | 41 | 5.09 | ROL.L# | 42 | -53 | -3 | 13 | 3.04 |
| PUT.L | 3093 | -30 | -9 | 7 | 4.18 |  |  |  |  |  |  |
| INS.L | 2692 | -34 | -4 | 10 | 4.33 |  |  |  |  |  |  |
| PreCG.L | 956 | -40 | -3 | 29 | 4.05 |  |  |  |  |  |  |
| ROL.L | 481 | -39 | -29 | 13 | 3.57 |  |  |  |  |  |  |
| IFGo.L | 439 | -39 | 7 | 19 | 3.69 |  |  |  |  |  |  |
| STG.L | 288 | -39 | -30 | 12 | 3.57 |  |  |  |  |  |  |
| PoCG.L | 214 | -36 | -13 | 39 | 3.54 |  |  |  |  |  |  |
| SMG.L | 203 | -48 | -36 | 30 | 3.42 |  |  |  |  |  |  |
| IFGtriang.L | 167 | -36 | 11 | 24 | 2.79 |  |  |  |  |  |  |
| MFG.L | 164 | -30 | 36 | 18 | 3.41 |  |  |  |  |  |  |
| HES.L | 150 | -36 | -29 | 15 | 3.52 |  |  |  |  |  |  |
| PAL.L | 59 | -20 | 1 | 8 | 3.35 |  |  |  |  |  |  |
| MTG.L | 33 | -47 | -41 | 8 | 3.57 |  |  |  |  |  |  |
| CAU.L | 31 | -21 | -13 | 23 | 2.81 |  |  |  |  |  |  |

|  |  |  |  |  |  |  |  |  |  |  |
| --- | --- | --- | --- | --- | --- | --- | --- | --- | --- | --- |
| AMYG.L | 10 | -29 | 4 | -17 | 3.06 |  |  |  |  |  |
| **<u>Overlap (Language Sequence ∩ Tool-use Sequence)</u>** |  |  |  |  |  |  |  |  |  |  |
| **Cluster1** | 13996 | -31 | 4 | -15 |  | **Cluster2** | 354 | -45 | -44 | 4 |
| WM | 10593 | -31 | 4 | -15 |  | WM | 299 | -45 | -44 | 4 |
| INS.L | 1598 | -36 | 4 | -13 |  | MTG.L | 33 | -48 | -44 | 5 |
| PUT.L | 372 | -31 | -5 | -7 |  | STG.L | 22 | -42 | -41 | 11 |
| ROL.L | 360 | -42 | -6 | 10 |  |  |  |  |  |  |
| STG.L | 249 | -40 | -4 | -10 |  | **Cluster3** | 45 | -28 | 6 | 5 |
| PreCG.L | 213 | -48 | -1 | 19 |  | PUT.L | 41 | -28 | 6 | 5 |
| IFGo.L | 205 | -39 | 11 | 16 |  | WM | 4 | -32 | 7 | 5 |
| SMG.L | 203 | -44 | -38 | 25 |  |  |  |  |  |  |
| PoCG.L | 114 | -49 | -2 | 18 |  | **Cluster4** | 43 | -22 | 35 | 14 |
| HES.L | 87 | -36 | -22 | 5 |  | WM | 43 | -22 | 35 | 14 |
| AMYG.L | 2 | -28 | 3 | -14 |  |  |  |  |  |  |

### Shared putaminal lesion patterns underlie goal-dependent impairments in language and tool use

To identify a shared organizing computation beyond the mere presence of sequences, we quantified goal dependency—the extent to which the constituent elements within a sequence are selected and structured according to the requirements of the intended outcome—across the two domains (Fig. 3A). In language, goal dependency was indexed by the proportion of well-formed sentences produced in the picture-description task, as structurally well-formed sentences necessarily preserve the goal-directed syntactic dependency constraints required for sentence construction, irrespective of semantic accuracy or lexical appropriateness (Ash et al., 2013; Harris et al., 2019; Rochon et al., 2000). In tool use, it was indexed by the expert-rated accuracy of functional tool grasp in the tool-use task, which captured whether the initial grasp configuration was selected according to the functional (goal) requirements rather than the morphological affordances of the tool alone. These behavioral indices of goal dependency correlated across patients (*r* = 0.38, *p* < 0.001). Lesion proportion within ten out of eleven VLSM-defined overlapping left-lateralized regions, notably the putamen and precentral gyrus, predicted the severity of the deficit in goal dependency measures in both language and tool use (FDR *p*s < 0.01; Fig. S2B).

**Figure 3.**
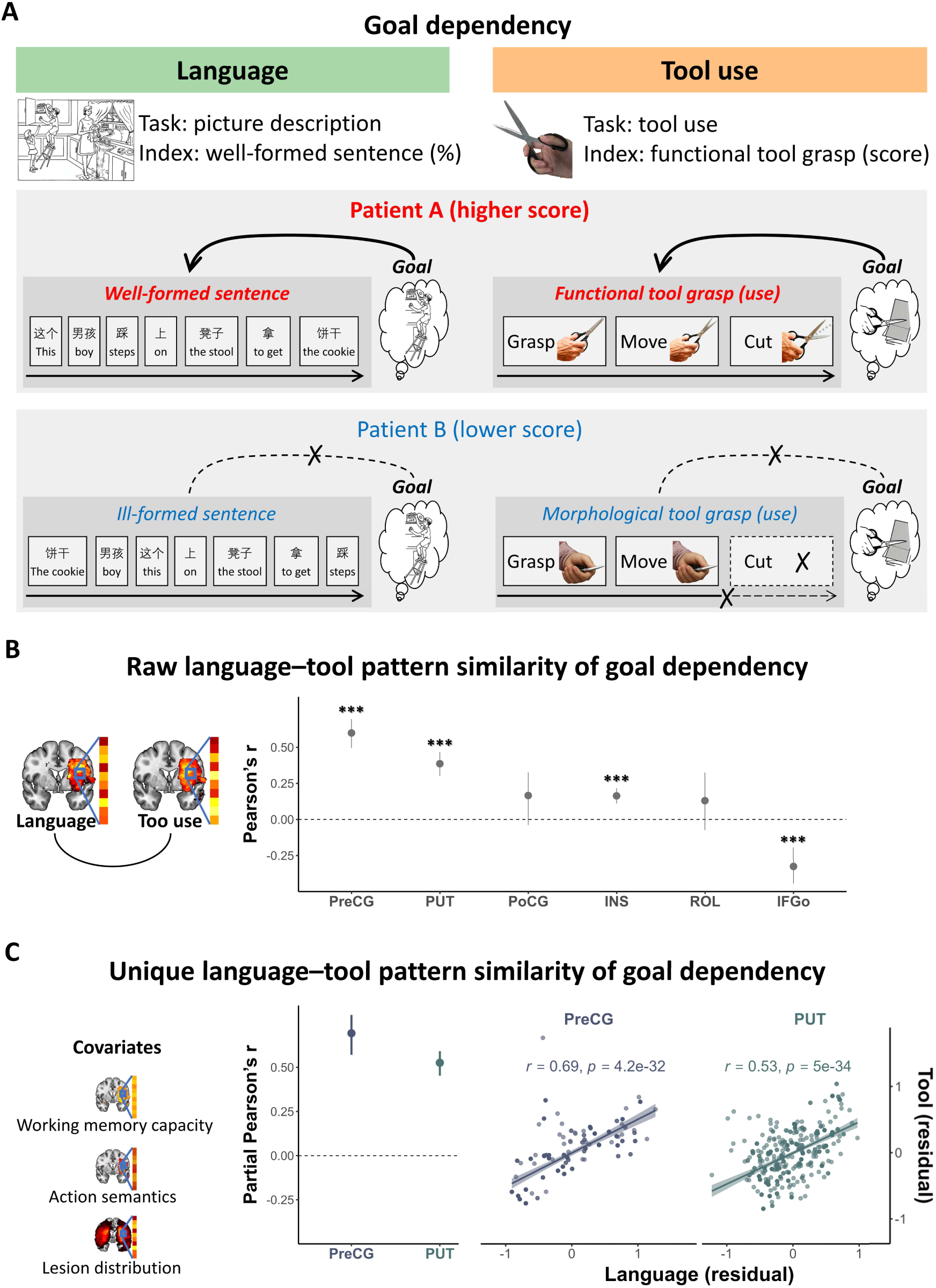
Study 1: Shared computation of goal dependency in language and tool use. ***A***, Behavioral manifestations and operationalization of goal dependency. Higher scores reflect the production of well-formed sentences in Cookie Theft picture description task for language, and using functional (goal-dependent) tool grasps for tool use. ***B***, Raw language–tool pattern similarity of goal-dependency. Voxel-wise Pearson’s correlations between VLSM t-scores of language goal-dependency and tool goal-dependency were significant in regions including the left putamen and left precentral gyrus. ROIs with sufficient voxel overlap (≥ 30 voxels) were selected a priori to ensure stable correlation estimates, resulting in the six ROIs shown here. Dots represent correlation coefficients and bars indicate 95% confidence intervals; ***FDR-corrected *p* < 0.001. ***C***, Unique language–tool pattern similarity of goal-dependency. Voxel-wise partial correlations remained selectively significant in the putamen and precentral gyrus after controlling for working-memory capacity, action semantics, and lesion distribution. The scatter plots display residual VLSM t-scores for goal-dependent sequence integrity of language and tool-use, computed after removing variance explained by these covariates. Each dot represents a voxel; regression lines show the partial Pearson’s correlation between language residuals and tool residuals, with shaded areas denoting 95% confidence intervals. Darker dots reflect a greater number of voxels occupying identical residual-value coordinates.

As regional overlap could reflect functionally distinct but spatially adjacent subregions, we examined cross-domain voxel-wise lesion-pattern similarity (i.e., multi-voxel pattern analysis of lesion effects, MVPA; Fig. 3B). VLSM-t maps associated with language- and tool-related goal-dependency deficits were spatially aligned in four regions, especially the putamen (*r* = 0.39, *p* < 0.001) and precentral gyrus (*r* = 0.60, *p* < 0.001). We further considered potential variables that are broadly shared across tasks and yet are not specific to these two domains: both language and tool-use tasks require the maintenance of sequential elements in working memory space; sentential stimuli often entail verbs related to actions that are also implicated in tool tasks. We thus tested whether the observed pattern similarity could be accounted for by working-memory capacity, action verb production (measured in the Cookie Theft task), or additionally by similar lesion distributions. These associations persisted only in the putamen and the precentral gyrus after controlling for working-memory capacity, action semantics, and lesion distribution (*r*s > 0.53; FDR *p*s < 0.001; Fig. 3C).

Specifically, patients with similar lesion patterns in the putamen or precentral gyrus showed parallel difficulties in constructing dependency relations across words in sentences and across action components in tool use. Although both regions contributed to performance in the current lesion analyses, only the putamen showed cross-study voxel-wise correspondence with the functional data (see below; Fig. S5). Together with the prior exclusion of domain-specific factors in the VLSM analyses (lexical semantics and action imitation), this pattern indicates that the shared neural architecture cannot be explained by shared non-structural, goal-independent influences, lesion distribution, or domain-specific factors, but instead reflects shared goal-dependency-based structure-building demands.

### Early language experience modulates tool-use behavior

Having established the causal necessity of the putamen for goal dependency processing across the language and tool domains in Study 1, we next asked whether early language experience developmentally shapes tool-use behavior in Study 2. We examined 40 congenitally deaf signers comprising two groups differing in the age of first language (sign language) exposure (Fig. 1, right panel; “native” signers exposed from birth; “delayed” signers exposed from a mean age of 7.42 years), with sex, age, education, and nonverbal intelligence quotient (IQ) well matched between groups (*p*s > 0.20). Tool use was indexed by the expert-rated correctness of goal-directed demonstrations, which were assessed using the same standardized procedure and scoring criteria as in Study 1 and applied to an expanded set of everyday tools. “Native” signers outperformed “delayed” signers on tool-use proficiency (*t*_(38)_ = 2.87, *p* = 0.007, *d* = 0.93; Fig. 4A).

**Figure 4.**
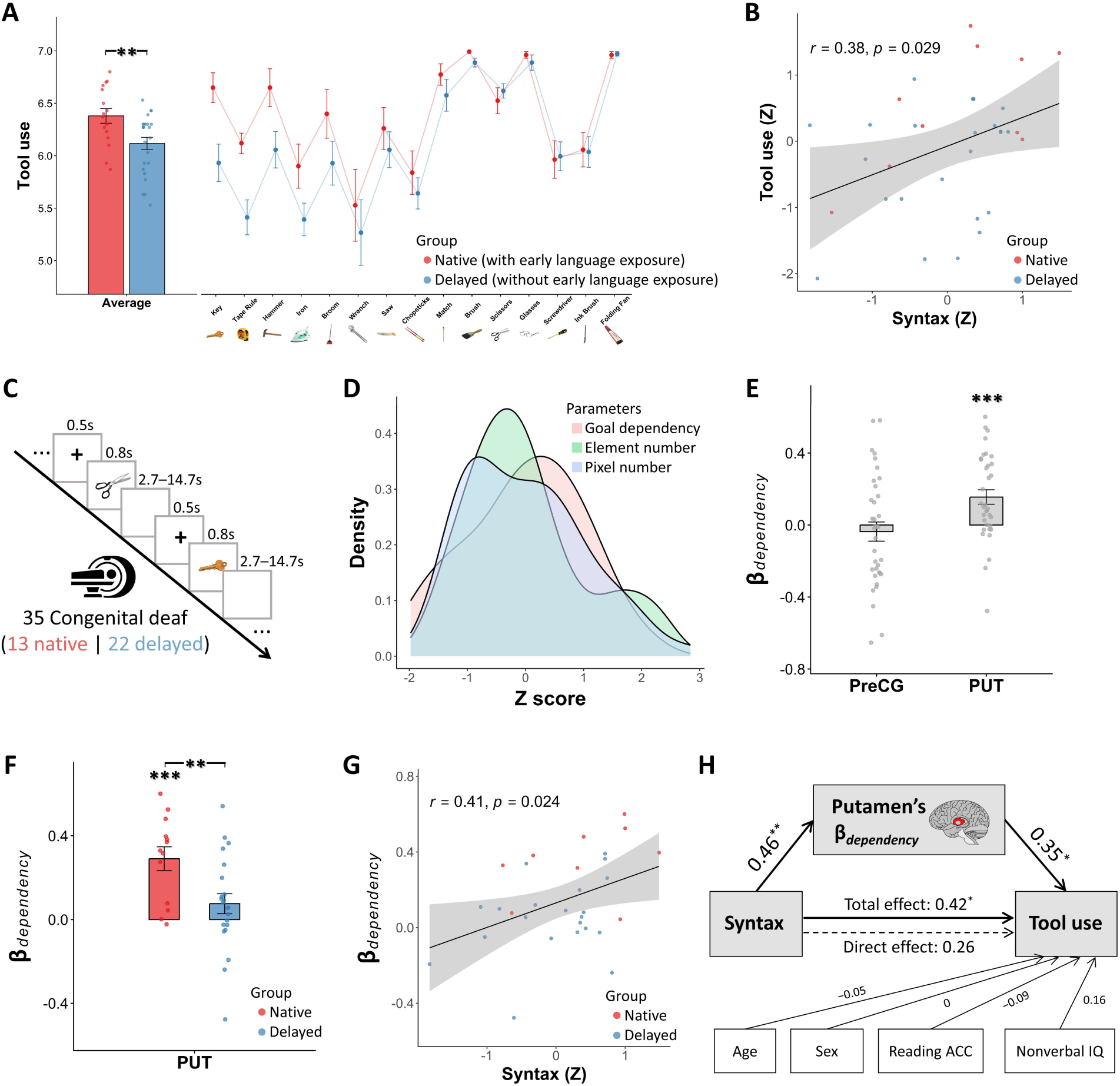
Study 2: Early syntactic experience modulates tool-use performance through putaminal encoding of goal dependency. ***A***, Compared with native signers, delayed signers showed lower tool-use performance. Each dot represents an individual deaf signer’s tool-use scores averaged across tool items (left panel) or group mean for each tool item (right panel). ***B***, Syntactic proficiency was positively correlated with tool-use performance across all the deaf signers. Each dot represents an individual deaf signer. ***C***, Schematic of the fMRI picture-sign naming paradigm. ***D***, Kernel density estimates show the distributions of goal dependency, element number, and pixel number across all artifacts, with values z-scored within each modulator. ***E***, ROI analyses revealed significant goal-dependency encoding in the putamen but not in the precentral gyrus. Each dot represents an individual deaf signer. ***F***, Native signers exhibited stronger putaminal goal-dependency modulation than delayed signers. Each dot represents an individual deaf signer. ***G***, Putaminal goal-dependency β values positively correlated with syntactic proficiency across all the deaf signers. Each dot represents an individual deaf signer. ***H***, The mediation analysis showed that the putaminal encoding of goal dependency fully mediated the syntax–tool-use relationship. General note: Error bars denote ± standard errors (SEs). Red dots denote native signers exposed to language from birth, whereas blue dots denote delayed signers exposed later in childhood (dots in D are all shown in grey). Regression lines in B and G depict Pearson’s correlations with shaded 95% confidence intervals. Significance: \**p* < 0.05, \*\**p* < 0.01 and \*\*\**p* < 0.001 (two-tailed; FDR-corrected).

This advantage cannot be attributed to sign-language–related manual experience: although delayed signers had more such experience than the healthy controls in Study 1, they did not exceed these controls on the tool-use items common to both studies (*t*_(65)_ = −2.60, *p* = 0.013, *d* = 0.69).

In parallel with the native–delayed group distinction in early language experience, their current syntactic competence was indexed by the expert-rated correctness of word-order in a sign-language Cookie Theft picture-description task. Across participants in Study 2, tool-use scores positively correlated with sign-language syntactic competence (*r* = 0.38, *p* = 0.03; Fig. 4B) but not with reading accuracy reflecting basic literacy skills (*r* = 0; Fig. S4). Together, these results indicate that early language experience, rather than manual experience or general literacy, supports the development of tool-use behavior.

### Putaminal encoding of goal dependency

We conducted parametric-modulation analyses to establish the role of the putamen in encoding goal dependency during tool cognition, which provided the basis for testing whether early linguistic experience modulates this encoding. These analyses quantified voxel-wise sensitivity to goal dependency from the fMRI responses to 95 pictures of objects (Fig. 4C). Although the participants did not perform an explicit tool use task in the scanner, viewing pictures of tools has been shown to automatically elicit brain activity underlying manipulation and functional knowledge representation of tools (Chao & Martin, 2000; Lewis, 2006). Accordingly, we interpret the present effects as reflecting the encoding of goal-dependent tool knowledge rather than online tool execution. The general linear model (GLM) included goal-dependency strength as the parametric modulator of interest, quantifying the degree to which action elements are organized into goal-dependency-structured representations. To control for potential confounds in picture viewing, the GLM additionally included the element number (indexing the working-memory load, associated with the number of action elements an object typically engages) and pixel number (indexing low-level visual complexity) as control modulators (distributions shown in Fig. 4D), with no detectable collinearity (Table S2). Orthogonalization was disabled to ensure that the observed effects of goal-dependency strength were not driven by order-dependent variance allocation among modulators. The goal-dependency strength and element number were obtained from independent behavioral ratings (N = 30 per measure), both of which showed high inter-rater reliability (*ICC*s > 0.9), whereas pixel number was computed from binarized object masks. Analyses focused on patient-derived ROIs (from Study 1) in the putamen and precentral gyrus.

Only the left putamen showed significant positive modulation (β) by the goal-dependency strength of objects (*t*_(34)_ = 3.84, *p* = 0.001, *d* = 0.65; Fig. 4E); no significant effect was observed in the precentral gyrus (*p* = 0.50, Fig. 4E). The ROI analysis confirmed stronger encoding of the goal-dependency strength than element number in the left putamen (*t*_(34)_ = 2.4, *p* = 0.02, *d* = 0.58).

These results further identify the putamen as a critical site involved in encoding the goal-dependency structure beyond the working-memory load or visual complexity of the pictures. Voxel-wise correlation between Study 2’s functional β-map for goal-dependency encoding and Study 1’s lesion–symptom t-map for tool’s goal-dependency measure revealed a convergent spatial organization in the left putamen (*r* = 0.69, *p* < 0.001; Fig. S5), confirming the cross-method convergence of goal-dependency-structured representations.

### Putaminal encoding mediates the syntax–tool interaction

Early syntactic experience selectively enhanced goal-dependency encoding in the left putamen (native vs. delayed signers: *t*_(33)_ = 2.82, *p* = 0.008; Fig. 4F). We also examined all other goal-dependency-sensitive regions identified at the whole-brain level (Fig. S6A). No significant differences between native and delayed signers were observed in any region outside the left putamen (*p*s > 0.18; Fig. S6B), indicating a putamen-specific effect. Across all the signers, syntactic competence significantly correlated with the strength of putaminal goal-dependency encoding (*r* = 0.41, *p* = 0.02; Fig. 4G), whereas reading accuracy did not (p = 0.98; Fig. S7B). Putaminal goal-dependency encoding, in turn, predicted tool-use performance (*r* = 0.42, *p* = 0.01; Fig. S7A). The left putamen thus may serve as the interface where linguistic experience tunes dependency-structured representations relevant for tool cognition.

A structural-equation model corroborated this mediation (Fig. 4H; controlling for age, sex, reading accuracy, and nonverbal IQ): syntax predicted putaminal goal-dependency encoding (β = 0.46, *p* = 0.004), which in turn predicted tool-use performance (β = 0.35, *p* = 0.03), while the direct syntax–tool path became nonsignificant (β = 0.26, *p* = 0.12). The indirect effect (β = 0.16) accounted for 38.2% of the total association between syntax and tool use and was significant based on bias-corrected bootstrap analyses (5,000 resamples; 95% confidence interval [CI] = [0.01, 0.48]). Although a mediation analysis alone cannot establish temporal causality, the observed pattern supports a developmental account in which early syntactic competence enhances adult tool cognition through its effect on putaminal encoding of goal dependency.

## Discussion

Across convergent methods—lesion mapping, developmental contrasts, and functional neuroimaging—we identify the putamen as a critical locus involved in regulating goal-dependent sequencing in both language and tool use (Fig. 5). Damage to this region selectively impairs the sequence structures during syntactic comprehension and functional tool use. Putaminal activity tracks the strength of goal-dependency, and its encoding mediates the influence of syntactic competence on tool-use performance. Together, these findings support a domain-general neural mechanism that regulates how serial events are organized to achieve anticipated outcomes, linking symbolic and action-based cognition through a shared control operation rather than shared representational content.

**Figure 5.**
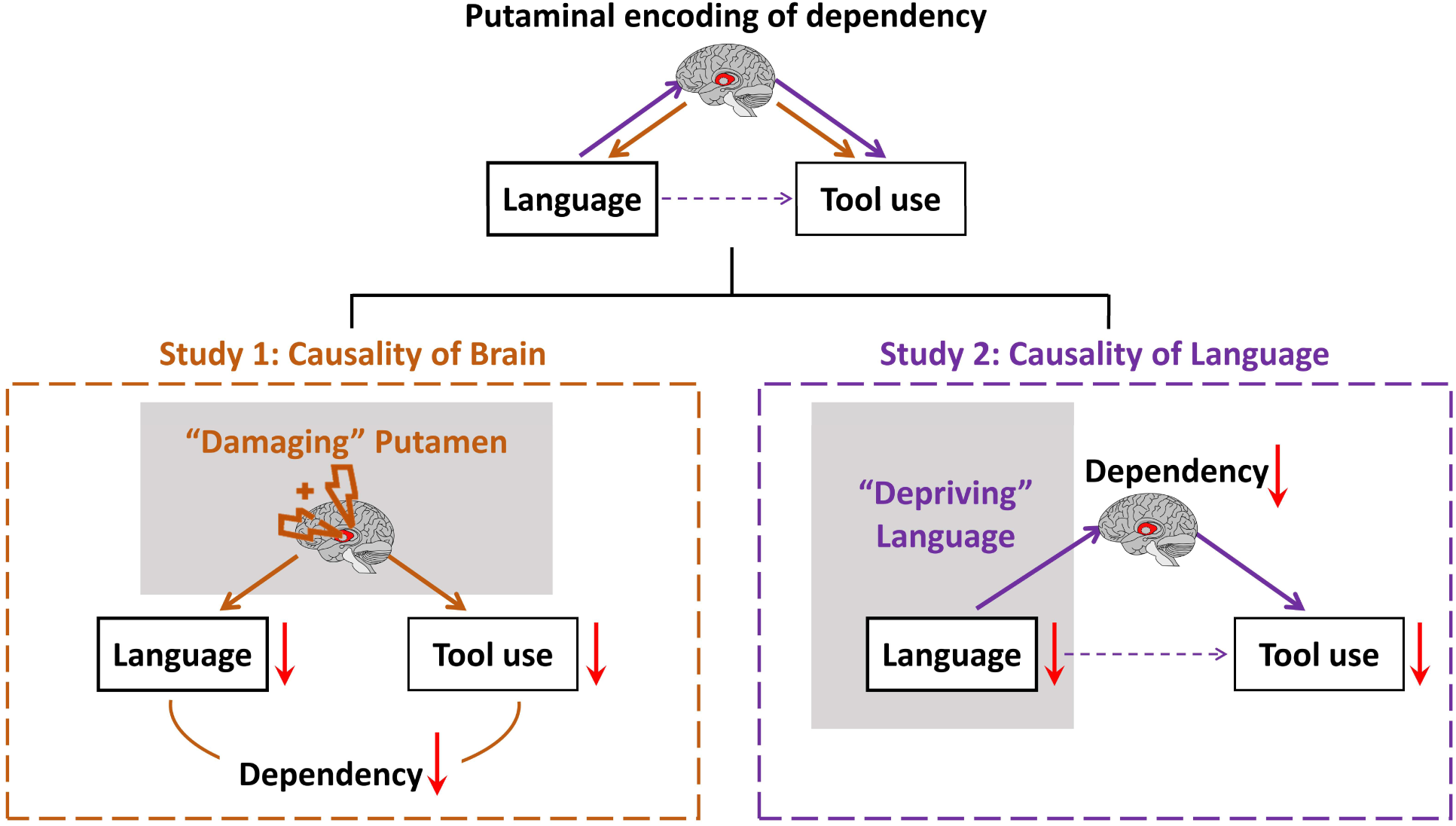
Summary of neural causal pathways linking language and tool use through the putaminal encoding of goal dependency. The schematic integrates findings from both studies to show that a shared computation—goal dependency encoded in the putamen—provides a common neural substrate for language and tool use. In Study 1, focal lesions to the putamen produced parallel impairments in goal-dependency of language and tool use, establishing lesion-based neural necessity. In Study 2, deprived language experience weakens the putaminal encoding of goal dependency and thereby reduces tool-use proficiency, revealing a developmental necessity through which language experience shapes this neural computation and its behavioral consequences.

Goal-dependent sequences are central to both language and tool use because they allow sequences to be organized and guided by distal outcomes that are not locally available at the time of early decisions, enabling abstract goals to constrain the selection and ordering of intermediate elements. This operation is unnecessary for simple reactive or habitual sequences, and is more relaxed in larger-scale action events, where different orderings of some intermediate elements can be treated as equivalent, but is particularly essential in forming sentences and tool-use action sequences (Coopmans et al., 2023; Greenfield & Westerman, 1978). By isolating this property, we identify a minimal computational demand that distinguishes language and tool use from broader types of sequential behavior.

Previous studies have reported overlapping activations for language and tool use in frontoparietal and basal ganglia networks, as well as cross-domain representational similarity during tool planning and syntactic processing (Higuchi et al., 2009; Thibault et al., 2021; Wen et al., 2024). However, these findings were correlational and did not identify the specific computational property underlying the observed overlap. By combining voxel-based lesion–symptom mapping with an explicitly specified, quantitative measure of goal-dependency, we demonstrate that lesions involving the putamen predict deficits in processing goal dependencies across both domains. This convergence is not explained by lexical semantics, motor execution, imitation, or working memory capacity, but instead reflects a shared operation that integrates future goal requirements into ongoing sequential choices.

We interpret the putamen as a key subcortical node within cortico–basal ganglia control loops that regulate the selection, maintenance, and updating of structured representations over time (Frank et al., 2001; Hazy et al., 2007; O’Reilly & Frank, 2006). Basal ganglia circuits are well established as mediators of sequential control through dopaminergic modulation and recurrent interactions with the cortex, supporting the stabilization of task-relevant information and the gating of alternative actions or representations (Alexander et al., 1986; Graybiel, 1998; Haber, 2016; Jin et al., 2014; Nicola et al., 2000). Deciding when to maintain, update, or release a dependency—an operation extensively characterized in motor sequence learning (Jin et al., 2014; Jin & Costa, 2010, 2015; Yang et al., 2025)—may generalize to syntactic and tool-related cognition without requiring the basal ganglia to encode domain-specific content. Instead, cortical regions provide representational specificity (words, gestures, and tool functions), while putamen-centered circuits regulate goal-dependency over these representations in a domain-general manner (Collins & Frank, 2013; Kaas & Stepniewska, 2023; Seger, 2008; Ullman, 2001, 2004).

Evidence from congenitally deaf adults further indicates that this goal-dependency regulation, while rooted in evolutionarily conserved sequencing circuitry, depends on early linguistic experience for full developmental calibration. Native signers—who were exposed to structured language from birth—outperformed delayed signers in tool-use tasks requiring goal-dependency integration and showed stronger putaminal sensitivity to goal-dependency strength. It is important to note that this advantage is not attributable to signing per se, as delayed signers, despite having richer signing experience than hearing controls assessed in Study 1, showed degraded tool-use performance. Mediation analyses indicated that syntactic competence influenced tool-use performance through its effect on putaminal encoding, suggesting that early language acquisition scaffolds the tuning of cortico–basal ganglia control mechanisms. Whether extended experience with tool use or other forms of complex action planning can similarly calibrate this system during development remains an open question.

These findings refine earlier proposals of a shared “grammar of action and language” (Coopmans et al., 2023; Greenfield, 1991; Greenfield & Westerman, 1978; Moro, 2014; Pastra & Aloimonos, 2012; Pulvermüller, 2014; Rizzolatti & Arbib, 1998; Thibault et al., 2021) by identifying a specific, quantifiable computational operation that unites the two domains. By isolating goal-dependency as a formal variable, we show that the putamen is a critical subcortical structure supporting linguistic and tool-use contexts. The conserved sequencing mechanisms of basal ganglia—linking sequential events into predictive units (Jin & Costa, 2010; Yang et al., 2025)—thus provide a substrate for goal-dependent sequence regulation that, in humans, is shaped and stabilized through language experience. Through evolutionary reuse and developmental calibration, this mechanism may underlie the human capacity to construct complex structures from serial experience, whether symbolic or motoric.

## Methods

### Study 1

#### Participants

The behavioral and neuroimaging data analyzed in Study 1 were collected from individuals with acquired brain damage and healthy controls, drawn from a large cohort established by our group (Bi et al., 2015; Fang et al., 2018; Han et al., 2013). A total of 100 brain-injured patients (20 females and 80 males), who were recruited from the China Rehabilitation Research Center, underwent structural MRI and completed behavioral assessments of the current study. The patient group had a mean age of 45.71 years (SD = 13.49, range = 19–76) and a mean educational attainment of 12.89 years (SD = 3.52, range = 2–22). The inclusion criteria were as follows: no history of prior brain injury; no comorbid neurological or psychiatric conditions (e.g., alcohol dependence or major depressive disorder); at least one-month post-onset (mean = 6 months, SD = 11, range = 1–85); and able to follow task instructions. Among these 100 patients, 54 had ischemic stroke, 30 had hemorrhagic stroke, 15 sustained traumatic brain injury, and 1 had toxic brain injury. The control group consisted of 43 healthy adults (18 females and 25 males) with no history of psychiatric or neurological disease. The controls had a mean age of 50.23 years (SD = 10.63, range = 26–72) and a mean of 13.79 years of education (SD = 3.70, range = 9–22).

The patient and control groups were comparable on key demographic variables, including age (*t*_(141)_ = −1.95, *p* = 0.06, Cohen’s *d* = 0.36) and years of education (*t*_(141)_ = −1.38, *p* = 0.17, Cohen’s *d* = 0.25). In contrast, and as expected, patients showed a significantly lower general cognitive status as indexed by the total score on the Chinese version of Mini-Mental State Examination (MMSE; Folstein et al., 1975) (*M*_patients_ = 21.77, *SD* = 7.77, *M*_controls_ = 28.70, *SD* = 1.12, *t*_(141)_ = −8.71, *p* < 0.001, Cohen’s *d* = −1.25).

All participants were native Chinese speakers and had provided written informed consent for their participation. All procedures were approved by the Institutional Review Board of the State Key Laboratory of Cognitive Neuroscience and Learning at Beijing Normal University.

#### Behavioral measurements of language and tool-use sequence

##### Sentence comprehension task

All the participants completed a sentence-picture matching task. In each trial, participants listened to a simple, complete spoken Mandarin sentence presented by the experimenter, while simultaneously viewing a sheet of paper displaying two vertically arranged pictures. One picture correctly depicted the content of the sentence, whereas the other was semantically incongruent. The participants were instructed to identify which picture matched the meaning of the sentence by pointing or giving a verbal response. The task included eight sentences, with the sentence order fixed across participants. The task accuracy was used for further analyses.

##### Tool use task

All the participants completed a tool-use task. In each trial, the experimenter handed a tool to the participant, who was asked to demonstrate its typical real-world use. Ten common household tools were included. All the sessions were video-recorded using a Canon digital camera for subsequent offline scoring (total duration: 12.4 h; patients: 9.51 h; controls: 2.92 h).

Two trained raters, who were blinded to the hypotheses of this study, independently evaluated the correctness of each demonstration using a 7-point Likert scale (1 = completely incorrect, 7 = completely correct), following a detailed operationalized scoring manual adapted from our previous study (Bi et al., 2015). The item-specific criteria are provided in Supplementary Table S1. The tool-use performance of each participant was quantified as the averaged ratings across raters and items. Inter-rater reliability was high across participants (*r* = 0.87, *p* < 0.001).

##### Control tasks

We evaluated participants’ performance in the following two non-sequence control tasks, which were later regressed out from sentence comprehension and tool-use measurements, respectively, to target sequence processing in language and tool use.

###### Word–picture verification

This task was to evaluate word-level semantic processing, a non-sequence component of the sentence comprehension task. In each trial, a word was presented at the top of the computer screen with a picture displayed below. Participants judged whether the word and picture referred to the same object or concept. Thirty non-tool word-picture pairs were included: 10 animals, 10 fruits and vegetables, and 10 faces. Performance was scored as the number of correct responses (range: 0–30).

###### Non-tool action imitation

This task involved non-tool-related intransitive actions and was used to quantify the impairment in producing actions in general. Participants were asked to view and imitate ten actions presented in ten short videos. All imitation sessions were video-recorded for subsequent offline scoring (total duration: 17.2 h; patients: 12.95 h; controls: 4.21 h). The two trained raters that scored tool use, who were blind to the hypotheses of this study, independently scored each imitation. Scores were assigned on a 7-point Likert scale based on similarity to the model action, following the same rating procedures used for the tool-use task. Operational scoring criteria for each action are provided in Supplementary Table S1. Inter-rater reliability was excellent (*r* = 0.91, *p* < 0.001).

#### Behavioral measurements of goal dependency in language and tool use

##### Goal dependency in language

Goal dependency in language was quantified here as individuals’ ability to produce syntactically well-formed sentences in the Cookie Theft picture oral description task from the Boston Diagnostic Aphasia Examination (Goodglass et al., 2001). Recordings from seven patients were unavailable, leaving 93 patients for analysis. Speech was audio-recorded using a Sony digital recorder for subsequent transcription (total duration: 8.54 h; patients: 7.32 h; controls: 1.22 h). Two trained researchers transcribed all recordings verbatim, with ambiguous syllables rendered in pinyin. Because patients’ speech was often fragmented, the two researchers manually segmented the transcripts into sentences, defining each semantically complete unit as a single sentence. All segmentation decisions were resolved by consensus. Each sentence was then manually evaluated for syntactic completeness, with well-formed sentences scored as 1 and others as 0. The evaluation was restricted to sentence-level syntactic structure, without regard to semantic accuracy or lexical appropriateness. For each participant, the proportion of well-formed sentences was calculated and used as the behavioral index of goal dependency in language.

To validate this manual metric, we computed an automated, computational syntactic complexity index based on three incremental parsing strategies—top-down, bottom-up, and left-corner parsing (Bird et al., 2009). The analytical pipeline included Chinese word segmentation using Jieba (https://github.com/fxsjy/jieba) and syntactic parsing using the Stanford Parser in Penn Treebank style (Klein & Manning, 2003; https://nlp.stanford.edu/software/lex-parser.shtml). For each parsing strategy, the number of parsing operations required to build a full parse was counted per sentence and then averaged across sentences for each participant. This complexity index was strongly positively correlated with the proportion of well-formed sentences (*r* = 0.57, *p* < 0.001; Fig. S2A), supporting the validity of the manual, structure-based syntactic measure.

##### Goal dependency in tool use

Goal dependency in tool use was quantified by grasp behavior in the tool-use task, as the initial grasp is constrained by higher-order functional goal in successful tool use (Buxbaum et al., 2003; Garcea & Buxbaum, 2019; Rosenbaum et al., 1990, 2001). The scoring rubric for functional tool grasp was adapted from the grasp-related criteria in the standardized tool-use manual (Supplementary Table S1). To ensure that this index was not driven by generalized motor impairment, two complementary dimensions were scored: morphological tool grasp, which measured the extent to which the grip conformed to the tool’s physical form, and tool movement, which assessed whether the manipulation phase matched canonical usage. All three dimensions (functional tool grasp, morphological tool grasp, and tool movement) were scored on the same 7-point scale by two independent raters. Four tools—chopsticks, scissors, broom, and folding fan—were selected for these ratings, considering that they are operated unimanually and do not involve complex bimanual coordination, allowing quantification of functional grasp versus morphological grasp in a well-defined tool use sequence while minimizing interference from peripheral motor impairments, particularly in patients with unilateral hemiplegia. The inter-rater reliability was high for each measure (*r*s > 0.85, *p*s < 0.001) and rating scores of each measure were averaged across raters and then across tools. Patients with scores ≤ 5 on any of the three dimensions were classified as failing the motor-screening criterion and excluded, leaving 88 patients for analyses.

##### Control behavioral measurements

To rule out the effects of potential variables that are broadly shared across language and tool use tasks and yet are not specific to these two domains, we considered the following two control behavioral measurements as covariates in the partial correlation analyses testing voxel-wise pattern similarity of goal dependency in language and tool use (Fig. S3).

###### Verbal working memory capacity

As both language and tool use tasks require maintaining sequential elements in working memory space, to rule out the possibility that the observed voxel-wise pattern similarity between the two domains was attributed to domain-general working-memory capacity rather than goal dependency per se, we obtained all participants’ performance of word repetition with two load levels. In the low-load condition, participants heard a single common two-character Chinese word and repeated it aloud immediately; two trials were administered, yielding a total score ranging from 0 to 2. In the high-load condition, participants heard a sequence of three common two-character Chinese words, completed an unrelated distractor task, and then attempted to recall all three words; performance was scored as the number of correctly recalled words (0–3). To obtain a measure of working-memory capacity independent of basic perceptual or articulatory ability, we computed the participant-level residual from regressing high-load scores on low-load scores.

###### Action semantics

Considering that sentential stimuli often entail verbs related to actions that are also implicated in tool tasks, to rule out the possibility that the observed voxel-wise pattern similarity between language and tool-use goal dependency was driven by action-semantic processing, we derived an action-semantics index from the speech transcripts of the Cookie Theft description task. Two trained researchers annotated all produced verbs as action verbs or non–action verbs following standard Chinese lexical-semantic classifications of Modern Mandarin Chinese (Huang & Liao, 2017), resolving discrepancies through discussion. For each participant, the action-semantics score was computed as, for each sentence, the number of action verbs divided by the total number of words in that sentence, and then averaged across all sentences produced by the participant.

#### Structural MRI acquisition and preprocessing

Structural MRI data for the brain-injured patients were acquired at the China Rehabilitation Research Center using a 1.5 T GE Signa Excite scanner (GE Healthcare, Milwaukee, WI, USA). High-resolution T1-weighted images were acquired in the sagittal plane using a 3D acquisition sequence with the following parameters: matrix = 512 × 512; voxel size = 0.49 × 0.49 × 0.70 mm³; field of view (FOV) = 25 cm; repetition time (TR) = 12.26 ms; echo time (TE) = 4.2 ms; flip angle = 15°; bandwidth = 11.9 Hz/pixel; inversion time (TI) = 400 ms; 248 slices. Each participant underwent two T1 scans, which were subsequently aligned and averaged to improve image quality. In addition, T2-weighted FLAIR images were acquired in the axial plane (matrix = 512 × 512; FOV = 250 × 250 mm²; voxel size = 0.49 × 0.49 × 5 mm³; TR = 8002 ms; TE = 127.57 ms; TI = 2000 ms; flip angle = 90°; 28 slices; bandwidth = 15.63 Hz/pixel; acquisition time = 4 min 48 s). FLAIR images were used solely as visual references to assist the manual lesion identification and were not included in subsequent analyses.

Image preprocessing was carried out as described in previous studies (Bi et al., 2015; Han et al., 2013). For each patient, the two T1-weighted structural scans were aligned and averaged using trilinear interpolation in Statistical Parametric Mapping (SPM, version 5; Wellcome Department of Imaging Neuroscience, London, UK; RRID:SCR_007037; http://www.fil.ion.ucl.ac.uk/spm/software/spm5). T2-weighted FLAIR images were co-registered to the averaged T1 image and resampled into the same native space. Lesion boundaries were manually delineated on the averaged T1 image by two trained researchers, working slice by slice with visual reference to the co-registered T2 FLAIR image. All the lesion masks were subsequently reviewed and confirmed by an experienced radiologist. Each structural image was resampled to a voxel resolution of 1 × 1 × 1 mm³ and normalized to Talairach (TAL) space using BrainVoyager QX 2.0 (Brain Innovation, Maastricht, The Netherlands; RRID:SCR_013057; https://www.brainvoyager.com). Normalization was performed by manually identifying anatomical landmarks, including the anterior commissure, posterior commissure, and the most extreme anterior, posterior, superior, inferior, left, and right points of the brain. Affine transformation between the native and TAL spaces was estimated using the Advanced Normalization Tools (ANTs, version 2.3.1; RRID:SCR_004757; http://stnava.github.io/ANTs/), and applied to the lesion masks. Finally, all lesion masks were transformed into the Montreal Neurological Institute (MNI) space, and all analyses and coordinates are presented in MNI space.

#### Whole-brain VLSM analyses of language and tool-use sequence

Voxel-based lesion–symptom mapping (VLSM; Bates et al., 2003) was used to identify brain regions supporting language and tool sequence. The language sequence was indexed by the standardized residuals of sentence comprehension scores after regressing out word–picture verification accuracy. The tool sequence was indexed by the standardized residuals of tool-use scores after regressing out action imitation performance. VLSM was conducted separately for each residualized measure. Total lesion volume was included as a covariate, and analyses were restricted to voxels lesioned in at least 10 patients. For each voxel, independent-samples *t*-tests compared behavioral performance between patients with and without damage to that voxel, yielding voxel-wise t statistics reflecting the contribution of each voxel to the behavioral measure. Positive t-values indicate worse performance in the lesioned than intact group. Resulting whole-brain t-maps were corrected using the false discovery rate (FDR, *q* < 0.05; Fig. 2B). To identify regions critical to both language and tool sequence, the two thresholded t-score maps were overlaid. Voxels surviving both thresholds were binarized (1 = shared, 0 = non-shared) and intersected with the Automated Anatomical Labeling atlas (AAL; Tzourio-Mazoyer et al., 2002). Shared voxels were localized to 11 AAL regions in the left hemisphere, which served as regions of interest (ROIs) for further analyses.

#### Voxel-wise pattern similarity analyses of goal dependency in language and tool use

As a preliminary validation prior to voxel-wise pattern analyses, we first examined whether goal dependency in language and tool use showed convergent lesion–behavior relationships at the ROI level. For each patient, these two measures were separately correlated with lesion proportion within each of the 11 ROIs identified above. Pearson’s correlations were computed and FDR corrected across ROIs (*q* < 0.05). Ten ROIs exhibited significant negative correlations with both behavioral measures, motivating further fine-grained voxel-level analyses within these regions.

Although overlapping univariate effects at the regional level suggest that language and tool use identify common anatomical regions, such overlap alone does not establish a shared computation, as functionally distinct but spatially adjacent subregions could drive the two effects. To test whether goal dependency in language and tool use is supported by a shared voxel-level mechanism, we assessed the spatial correspondence between voxel-wise lesion–behavior associations (i.e., MVPA of lesion effects) for the two domains. That is, we computed the voxel-wise Pearson’s correlations between the raw, unthresholded VLSM t-maps for goal dependency of language and tool use. In the VLSM framework, positive t-values indicate voxels whose damage is associated with worse behavioral performance. Accordingly, voxel-wise correlations were restricted to voxels showing positive t-values in both maps. ROIs with at least 30 voxels (six regions shown in Fig. 3B) were included in this analysis to mitigate unstable correlation estimates driven by very small voxel counts. Within each ROI, voxel-wise Pearson’s correlations were computed across voxels between the two maps (FDR corrected, *q* < 0.05; Fig. 3B).

For ROIs showing significant voxel-wise pattern correlations, we further conducted voxel-wise partial correlation analyses to evaluate whether the observed alignment persisted after accounting for the potentially shared processes. These analyses controlled for the VLSM t-map of working-memory residuals, the VLSM t-map of the action-semantics index, and voxel-wise lesion counts (number of patients with damage to each voxel). Within each ROI, voxel-wise partial correlations were computed across voxels between the two residualized maps (FDR corrected, *q* < 0.05).

### Study 2

#### Participants

The behavioral and neuroimaging data analyzed in Study 2 were collected from 40 congenitally deaf signers with differing early language experiences: 16 “native” signers who acquired sign language from birth in deaf-parented families, and 24 “delayed” signers who were born to hearing parents and acquired their first language (sign language) later in childhood (with mean age of sign language acquisition of 7.42 years, SD = 2.06, range = 3–12). Participants were recruited through special education schools and community networks. All participants completed a background survey assessing hearing status, early language exposure, educational history, and nonverbal IQ measured using the nonverbal subtest of the Kaufman Brief Intelligence Test–Second Edition (KBIT-2; Kaufman, 2004). They had normal or corrected-to-normal vision, and reported no history of neurological or psychiatric disorders. Native signers (6 females and 10 males) had a mean age of 32.63 years (SD = 6.86, range = 21–45), completed an average of 15.13 years of formal schooling (SD = 2.39, range = 9–16). Delayed signers (14 females, 10 males) had a mean age of 31.71 years (SD = 4.99, range = 22–40), completed 15.83 years of formal schooling on average (SD = 0.82, range = 12–16). The two groups did not differ in age (*t*_(38)_ = 0.49, *p* = 0.63), sex distribution (χ²_(1)_ = 1.67, *p* = 0.20), years of education (*t*_(38)_ = 1.14, *p* = 0.27), and nonverbal IQ (*t*_(38)_ = 0.51, *p* = 0.61).

All participants provided written informed consent and received monetary compensation for their participation. All procedures were approved by the Institutional Review Board of the State Key Laboratory of Cognitive Neuroscience and Learning at Beijing Normal University.

#### Behavioral measurements

##### Tool-use task

All 40 deaf signers completed the tool-use task. The procedure was identical to that of Study 1. The current task included the same 10 tools used in Study 1, along with five additional items, resulting in a total of 15 tools. All the sessions were video-recorded using the same experimental setup as in Study 1 for subsequent offline scoring (total duration: 1.22 h).

Behavioral performance was scored using the same 7-point Likert scale, scoring manual, and item-level criteria as in Study 1 (Supplementary Table S1). Two independent raters, who were blind to the participants’ group status and the hypotheses of this study, scored all the items. Inter-rater reliability across participants was good (Pearson’s *r* = 0.73, *p* < 0.001). All analyses were based on the overall tool-use correctness scores averaged across raters and items.

##### Cookie Theft picture description task

Thirty-five deaf participants (including 13 native signers and 22 delayed signers) described the Cookie Theft picture (the same stimulus used in Study 1) in sign language (1 native signer and 1 delayed signer were excluded from analysis due to non-compliance with task instructions). Responses were video- and audio-recorded using a Canon camcorder for subsequent offline scoring (total duration: 1.28 h). The analysis focused on word order in sign language, a key syntactic dimension of signed utterances (Napoli & Sutton-Spence, 2014; Sandler & Lillo-Martin, 2006). Annotation and scoring procedures were carried out by three expert annotators using ELAN (version 6.6; RRID:SCR_014202), who were native deaf signers with formal linguistic training. Prior to formal annotation, the annotators jointly reviewed three example videos to establish shared criteria for sentence segmentation and then segmented the signed utterances from each participant. The resulting segmentations were jointly reviewed and reconciled through group discussion to produce a single consensus segmentation for each participant.

For scoring, the same annotators again used the three example videos to standardize the interpretation of the evaluation criteria. They then independently rated each segmented sentence produced by the 33 participants on a 7-point Likert scale (1 = highly unnatural word order, 7 = highly natural, 4 = neutral), focusing exclusively on the naturalness of word-order patterns, irrespective of semantic accuracy or relevance to the picture. For each annotator, scores were averaged across sentences and then standardized (z-scored) across all participants. Each participant’s final word-order score was computed as the mean of the three annotator-specific z-scores. The inter-rater reliability of the word-order ratings was good (*ICC*_(2,3)_ = 0.80, 95% CI [0.64, 0.89], *F*_(32, 64)_ = 4.85, *p* < 0.001).

##### Control behavioral measurement: Chinese sentence reading

All 40 deaf participants completed a Chinese reading task, which required semantic judgments of visually presented Chinese sentences. In each trial, either a complete Chinese sentence (e.g., “Cucumbers are spherical in shape”) or a string of non-linguistic symbols (e.g., “$$$$”) was presented at the center of the screen. When a sentence was presented, participants were instructed to judge whether its meaning was semantically correct, pressing the “Y” key with their right index finger for semantically correct sentences and the “U” key with their right middle finger for incorrect sentences. When a symbol string was presented, they pressed the space bar with their right thumb. The task included 50 trials of semantically correct sentences, 50 trials of semantically incorrect sentences, and 25 trials of symbol string. Each trial began with a 1000-ms fixation cross, followed by the stimulus, which remained on the screen until a response was made. The next trial started immediately after the keypress. The response accuracy was recorded. The task was programmed and implemented using PsychoPy (version 2021.2.3; RRID:SCR_006571; https://www.psychopy.org; Peirce et al., 2019).

##### fMRI picture sign-naming task

Thirty-five out of the 40 deaf participants (13 native signers and 22 delayed signers) completed an fMRI picture sign-naming task (Fig. 4C). The remaining five participants did not undergo MRI scanning due to contraindications (e.g., metal implants or tattoos) or time constraints. During scanning, participants were asked to produce the sign corresponding to each visually presented object picture with their left hand only to reduce motion artifacts.

The stimulus set consisted of 95 colored pictures of real-world objects presented on a white background, including 63 artifact items and 32 non-artifact animal items. Artifact items primarily included common manmade tools and utensils (kitchen implements, stationery, medical instruments, and personal accessories). All pictures were collected from publicly available online sources and standardized to 400 × 400 pixels (visual angle: 10.55° × 10.55°).

The task comprised six functional runs, each lasting 528 s. In each run, all 95 pictures were presented once in a pseudorandomized order. Each trial began with a 0.5-s fixation cross, followed by a 0.8-s picture presentation. Intertrial intervals varied from 2.7 to 14.7 s, with stimulus order and timing optimized using optseq2 (http://surfer.nmr.mgh.harvard.edu/optseq/). Each run was preceded and followed by a 10-s blank screen. The task was programmed and presented using E-Prime 2.0 (Version 2.0; Psychology Software Tools, Inc.; RRID:SCR_009567; https://pstnet.com/products/e-prime/).

#### Functional MRI acquisition and preprocessing

Functional and structural MRI data were acquired for the 35 deaf participants at the Neuroimaging Center of Beijing Normal University using a 3T Siemens Trio Tim scanner (Siemens, Erlangen, Germany). Functional images were acquired during the picture sign-naming task using a gradient-echo echo-planar imaging (EPI) sequence: 33 axial slices; TR = 2000 ms; TE = 30 ms; flip angle = 90°; matrix = 64 × 64; voxel size = 3 × 3 × 3.5 mm³; interslice gap = 0.7 mm. High-resolution structural images were collected using a sagittal 3D magnetization-prepared rapid gradient echo (MPRAGE) sequence (144 slices; TR = 2530 ms; TE = 3.39 ms; flip angle = 7°; matrix = 256 × 256; voxel size = 1.33 × 1 × 1.33 mm³).

Task-fMRI data were preprocessed using Statistical Parametric Mapping (SPM, version 12; RRID:SCR_007037; https://www.fil.ion.ucl.ac.uk/spm) implemented in MATLAB (MathWorks; RRID:SCR_001622; https://www.mathworks.com). For each functional run, the first five volumes were discarded to allow for signal equilibrium. The functional images were corrected for slice acquisition timing and realigned to the first volume of the first run using a six-parameter rigid-body transformation. Each participant’s T1-weighted anatomical image was coregistered to the mean functional image. Normalization parameters were estimated from the T1-weighted image using unified segmentation and applied to the functional images, which were then smoothed with a Gaussian kernel of 6 mm full-width at half-maximum (FWHM). Head motion parameters were extracted during realignment, and runs exceeding 3 mm of translation or 3° of rotation were excluded. Given the rarity of congenitally deaf signer populations, motion exclusion was applied at the run level rather than at the participant level. Participants with at least four usable runs were retained for analyses. Consequently, 30 participants retained all six runs, 3 retained five runs, and 2 retained four runs, yielding a total of 202 usable runs across 35 participants for subsequent analyses.

#### First-level parametric modulation analysis of goal dependency in tool processing

To identify brain regions sensitive to goal dependency, we employed a parametric-modulation approach in which three picture-wise features—goal dependency, element number, and pixel number—were entered as parametric regressors in the fMRI General Linear Model (GLM). Goal dependency and element number were obtained through online questionnaires administered on the Credamo platform (Credamo; Beijing Yishu Mofa Technology Co., Ltd; https://www.credamo.com/), each of which was rated by a separate group of 30 participants who did not participate in the fMRI experiment. The participants evaluated each picture on a 7-point Likert scale. For goal dependency, participants were asked to judge the extent to which the sub-actions required to interact with an object are interdependent and sequentially constrained (1 = very weak dependency; 7 = very strong dependency). For the element number, participants were asked to judge the number of simple sub-actions required to interact with the object (1 = very few; 7 = many). Ratings were averaged across raters to yield one score of goal dependency and one score of element number per picture, both of which showed inter-rater reliability (goal dependency: ICC_(2,30)_ = 0.96, 95% CI [0.94, 0.97], F_(94, 2726)_ = 22.07, *p* < 0.001; element number: ICC_(2,30)_ = 0.92, 95% CI [0.89, 0.94], F_(94, 2726)_ = 11.77, *p* < 0.001). The pixel number was computed to control for picture visual complexity following established procedures (Fan et al., 2021). Pictures were converted to grayscale; a binary object mask was generated by thresholding all pixels with grayscale values < 240. The total pixel number within the object mask was counted as a low-level visual-complexity measure. All three features were z-scored across pictures and entered as parametric modulators. Multicollinearity among the three features was assessed in SPSS Statistics (version 26; IBM Corp.; RRID:SCR_016479; https://www.ibm.com/analytics/spss-statistics-software). All variance inflation factors (VIFs) were less than 2.5, indicating no problematic collinearity (Table S2).

At the individual-subject level of parametric modulation, the GLM of each participant modeled artifact (n = 63) and animal (n = 32) pictures as separate conditions, with distinct onset regressors for each. The three parametric modulators were applied only to artifact trials and were entered simultaneously with the orthogonalization disabled (SPM: orth = 0), ensuring that shared variance was not redistributed based on regressor order. Onsets were time-locked to the picture presentation and convolved with the canonical hemodynamic response function. Six head-motion parameters were included as nuisance regressors for each run, and a 128-s high-pass filter was applied. Following prior work (Fan et al., 2021), we had no a priori rationale to expect negative effects of feature load; therefore, analyses focused on positive parametric effects (feature parameter estimate > 0).

#### Effects of language experience on the neural encoding of goal dependency

Neural encoding of goal dependency was first evaluated at the ROI level, followed by whole-brain analyses. The ROIs focused on the lesion-defined left putamen and left precentral gyrus ROIs in Study 1, as they demonstrated significant cross-domain voxel-wise lesion pattern similarity after controlling for potential confounding variables (see Results). These lesion-defined ROIs (1-mm isotropic) were resampled to the 3-mm isotropic to match the spatial resolution of the functional data in Study 2, using nearest-neighbor interpolation in DPABI (version 9.0; The R-fMRI Network; https://rfmri.org/dpabi) (Yan et al., 2016). Goal-dependency contrast (beta) estimates were then averaged across all voxels within each lesion-defined ROI for each participant. These ROI-level values were then tested against zero using one-sample t-tests across participants to evaluate whether a given ROI significantly encodes goal dependency. These values were compared between native and delayed deaf signers using two-sample t-tests (two-tailed) or correlated with the behavioral measures (e.g., the expert-rated syntactic competence, tool-use performance) using Pearson’s correlation.

To assess the neural encoding of goal dependency beyond the lesion-derived ROIs, whole-brain parametric-modulation analyses were conducted across all deaf signers. Whole-brain statistical maps were thresholded using voxel-wise FDR correction of q < 0.05, with an additional cluster-extent threshold of > 40 voxels. For each significant cluster, mean β values were extracted and compared between native and delayed signers using independent-sample t tests (two-tailed).

Finally, to test whether the putaminal encoding of goal dependency mediated the relationship between syntactic competence and tool use, we conducted structural equation modeling (SEM). In the SEM, syntactic competence was specified as the predictor, tool-use performance as the outcome, and putaminal goal-dependency β values as the mediator. Sex, age, Chinese reading accuracy and nonverbal IQ were included as covariates predicting tool-use performance. SEM was implemented in Amos 20.0 (IBM SPSS Amos; RRID:SCR_014498; https://www.ibm.com/analytics/amos) with the full-information maximum-likelihood estimation, according to standard procedures (Arbuckle, 2011). Path coefficients are reported as standardized estimates. Model fit was evaluated using standard indices of model fit, including the root-mean-square error of approximation (RMSEA). To obtain a robust test of the indirect (mediated) effect without relying on normality assumptions, bias-corrected bootstrap confidence intervals (5,000 resamples) were computed.

## Acknowledgments

This work was supported by National Natural Science Foundation of China (32595491 to Y.B.; 32171052 to X.W.), Brain Science and Brain-like Intelligence Technology—National Science and Technology Major Project 2021ZD0204100 (2021ZD0204104 to Y.B.), Major Project of National Social Science Foundation (24&ZD252 to Z.H.) and the China Postdoctoral Science Foundation (2024M760231 to H.W.). We thank Ziyi Xiong, Nai Ding, Hang Zhang, Wei Liu, Shufan Mao, Xitong Liang, Yuxi Chu for helpful discussions; Xuan Zheng and Xi Yu for guidance with performance measurements of deaf signers; Luping Song for assistance with data collection from patients; Shuyue Wang and Haoyang Chen for assistance with data collection from deaf signers; Chengcheng Wang for assistance with syntactic performances measurements of patients; Dingchen Zhang, Rui Feng, Miao Heng, Bing Chou, Yongsheng Wu for rating the performance of patients or deaf participants. We also thank all subjects who participated in our study.

## Author contributions

Y.B. conceived the study. Z.F., H.W., X.W. and Y.B. designed the experiment. Z.F., X.W. and Z.H. collected the data. Z.F. analyzed the data. Z.F., H.W. and Y.B. wrote the initial draft. Z.F., H.W., X.W. and Y.B reviewed and edited the Article.

---

**Figure S1.**
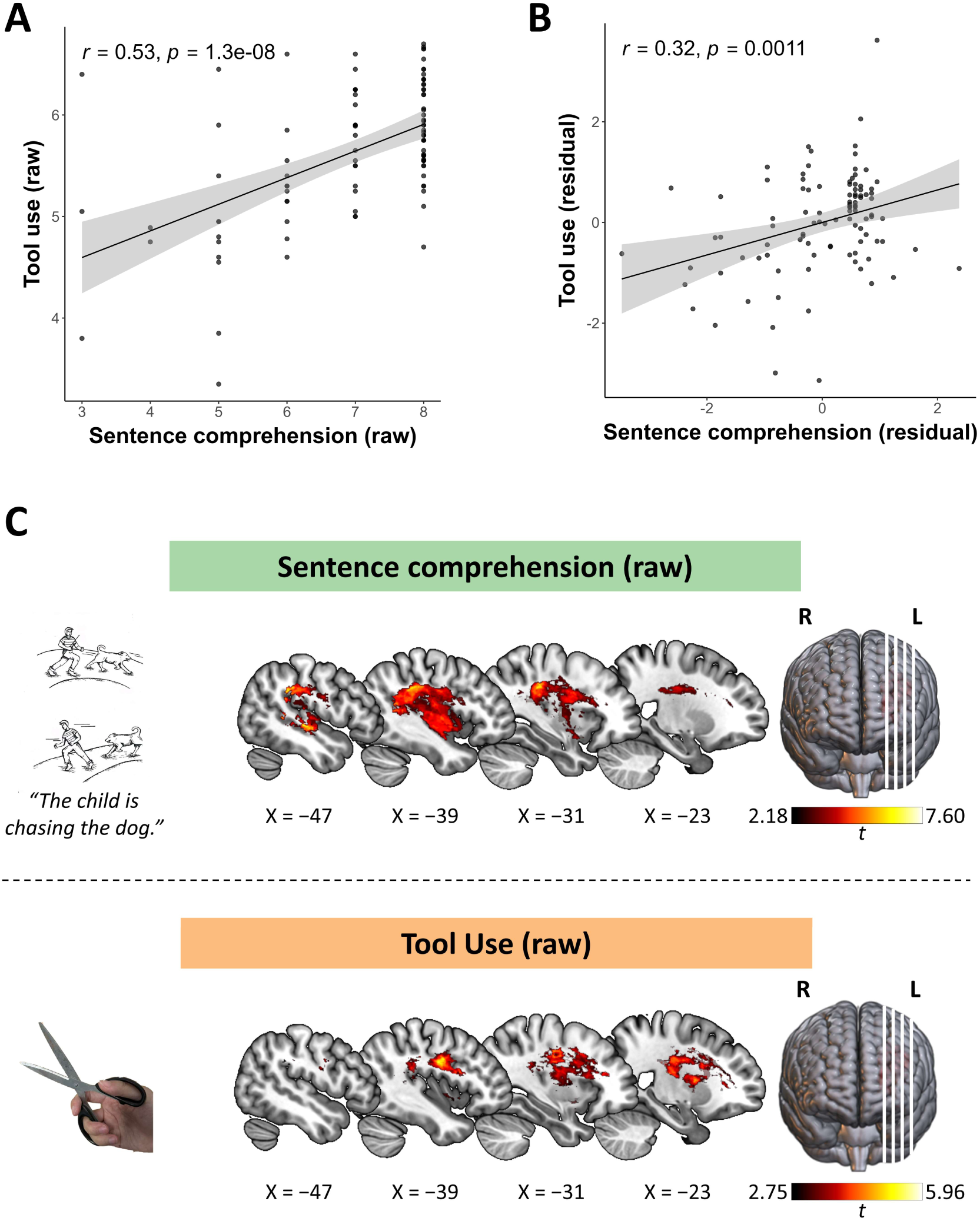
Study 1: Behavioral associations and lesion substrates for sentence comprehension and tool use. ***A***, Pearson’s correlation between raw sentence comprehension accuracy and raw tool-use performance across brain-injured patients. Each dot represents an individual patient. ***B***, Pearson’s correlation between residualized sentence comprehension (language sequence) and residualized tool-use performance (tool sequence) across brain-injured patients, obtained after regressing out word-level semantic processing and non-tool action imitation, respectively. ***C***, VLSM results for raw scores of sentence comprehension (top panel) and tool use (bottom panel) (voxel-wise FDR corrected, *q* < 0.05). The color bars show voxel-wise t-values. L denotes the left hemisphere, and R denotes the right hemisphere in all panels.

**Figure S2.**
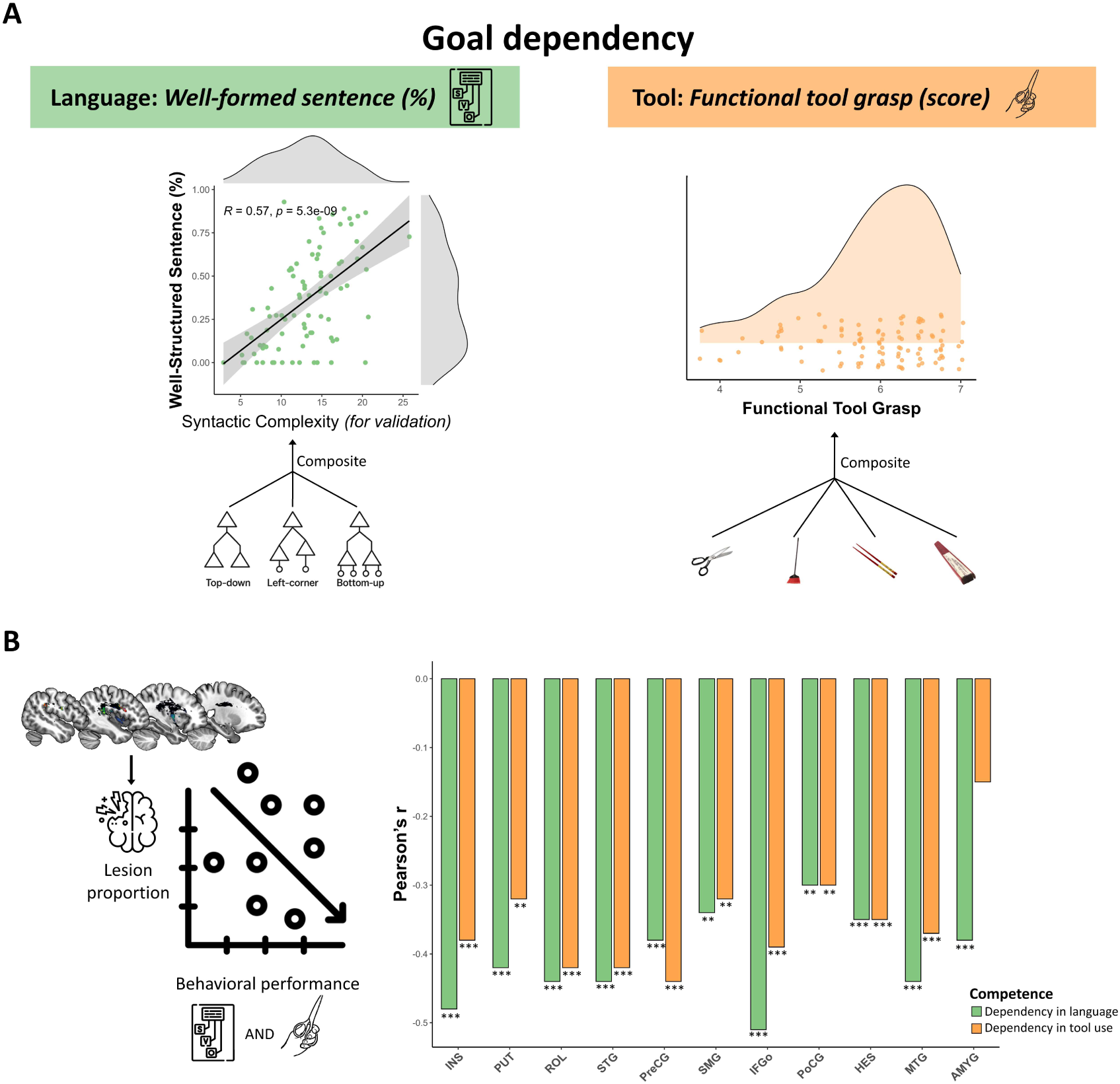
Behavioral distributions and lesion–behavior associations within overlapping lesion-defined ROIs. ***A***, Distributions of behavioral indices indexing goal-dependent sequence in language and tool use. The left panel shows the distribution of well-formed sentence production and its validation against syntactic complexity, which was computed using three parsing schemes (top-down, left-corner, and bottom-up). The right panel shows the distribution of functional tool grasp, computed from four tool types differing in action structure (scissors, broom, chopsticks, and folding fan). Each dot represents an individual patient in both panels. ***B***, ROI-wise associations between lesion proportion and behavioral performance across the 11 overlapping lesion-defined regions.The bars show Pearson’s r between the lesion extent and each behavioral index of goal-dependent sequence (language is shown in green; tool is shown in orange). FDR-corrected significance: *p* < 0.01** and *p* < 0.001***, two-tailed.

**Figure S3.**
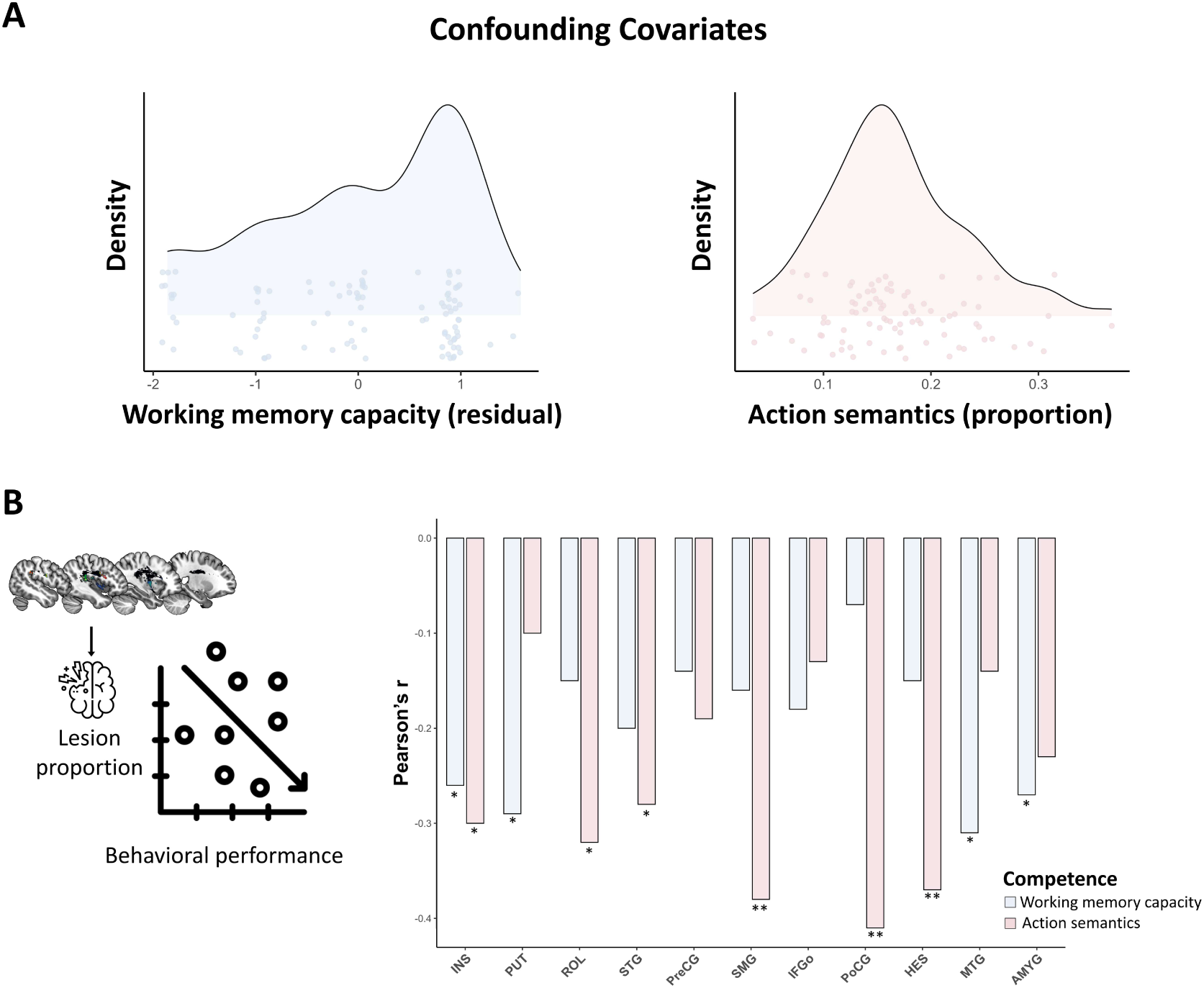
Confounding covariates: distributions, and ROI-wise lesion–deficit associations. ***A***, Distributions of the covariates. Kernel density plots with individual dots show working-memory capacity (left panel, light blue) and action semantics (right panel, light rose). Each dot represents an individual patient in both panels. ***B***, ROI-wise associations between lesion proportion and the covariates within 11 overlap-defined ROIs. The bars show Pearson’s r values for working memory capacity (light blue) and action semantics (light rose). Negative correlations indicate that a greater lesion extent predicts poorer performance. FDR-corrected significance: *p* < 0.05* and *p* < 0.01**, two-tailed.

**Figure S4.**
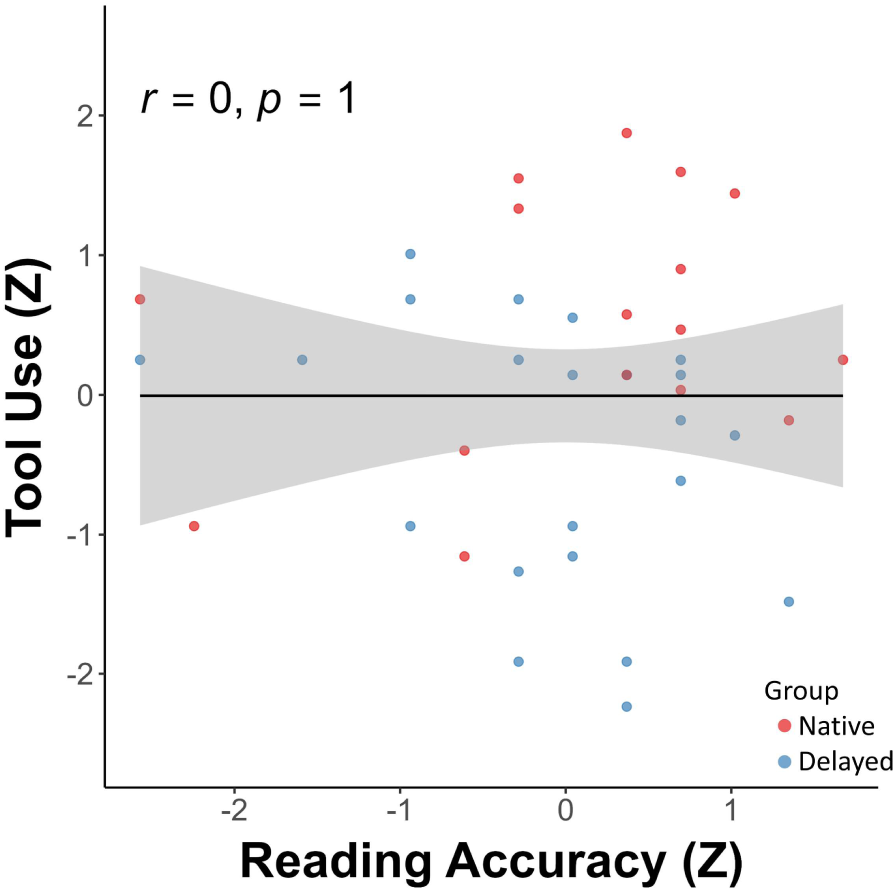
No association between Chinese reading accuracy and tool-use performance in deaf signers. Each dot represents an individual deaf signer (red = native; blue = delayed). The regression line shows the fitted relationship (*r* = 0, *p* = 1), with shaded areas denoting the 95% confidence interval.

**Figure S5.**
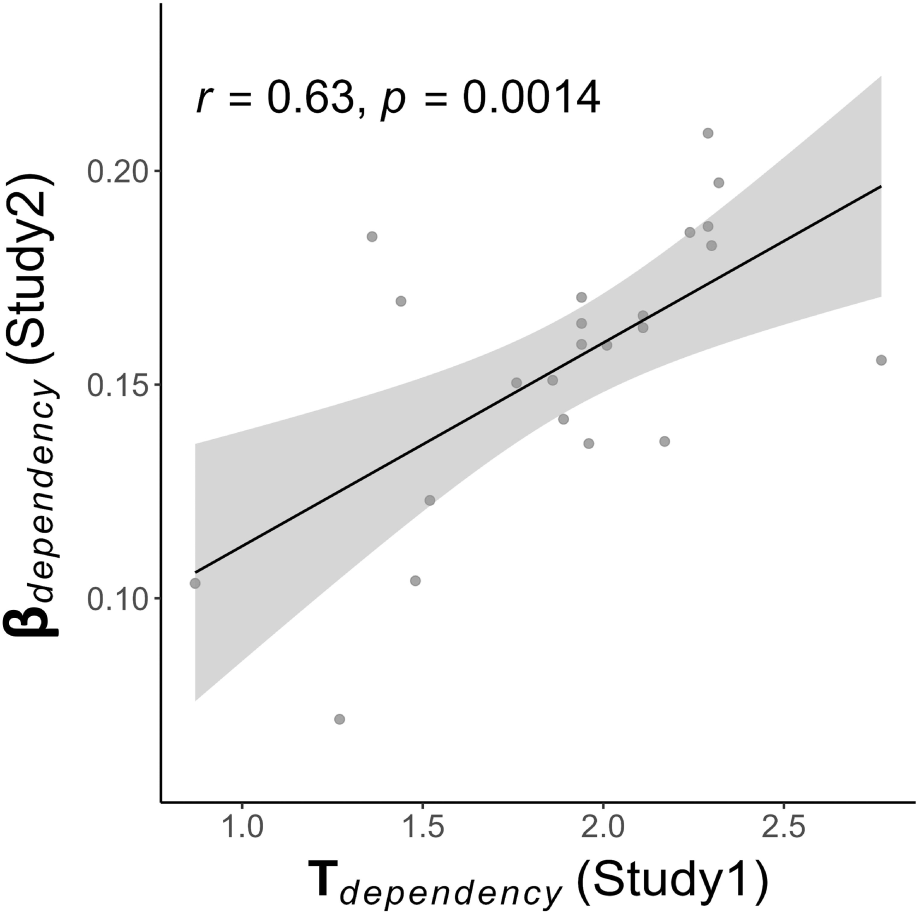
Cross-study voxel-wise correspondence of putaminal goal-dependency encoding. Voxel-level β values indexing artifact’s goal dependency in Study 2 were significantly correlated with VLSM t-scores for tool-related goal dependency in Study 1 (*r* = 0.63, *p* = 0.0014). The regression line depicts the fitted relationship, with shaded bands showing the 95% confidence interval. Each dot represents a single voxel within the left putamen ROI.

**Figure S6.**
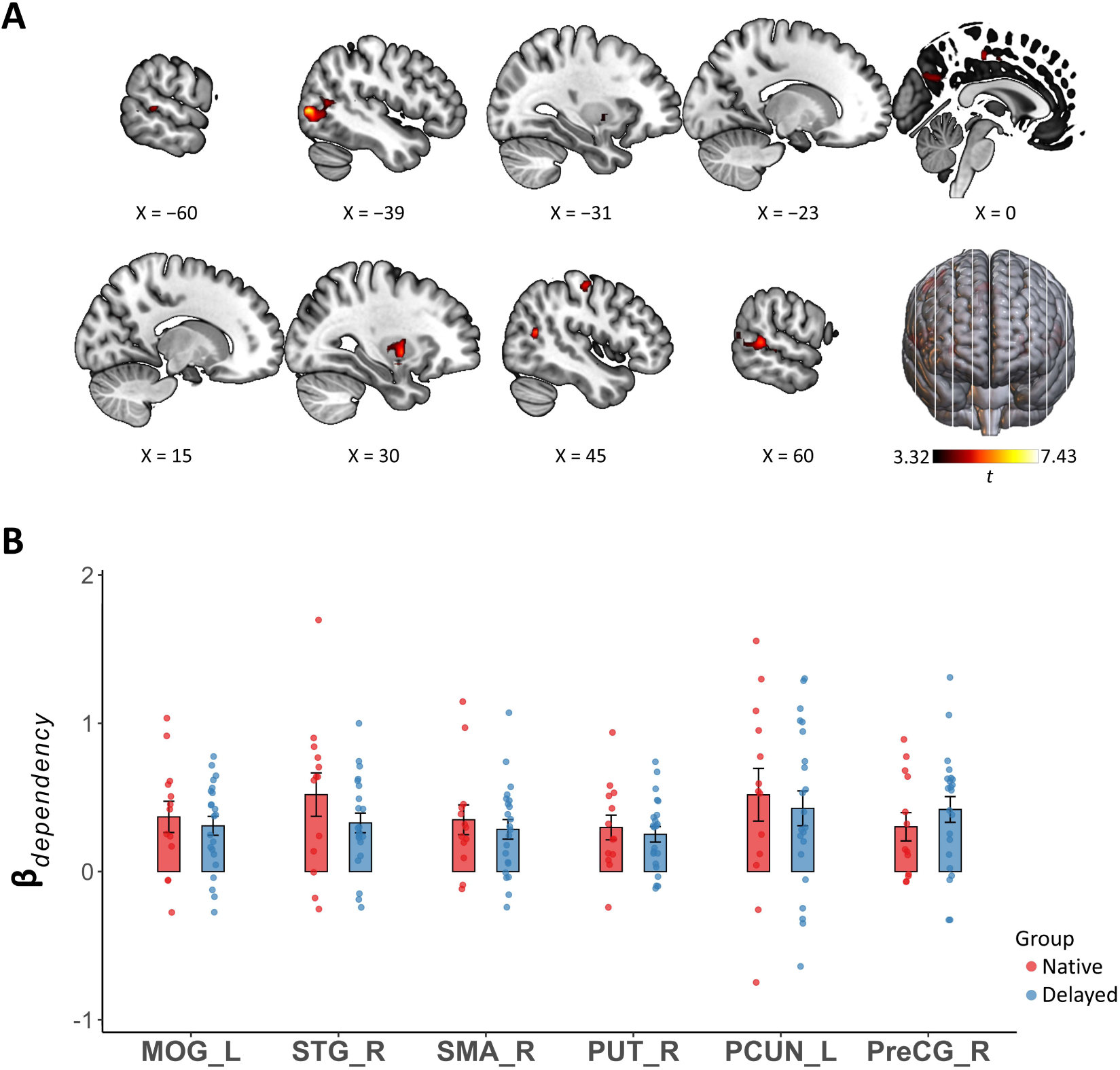
No significant group differences in goal dependency encoding across whole-brain dependency-sensitive clusters outside the left putamen. ***A***, Whole-brain parametric-modulation results for goal dependency (voxel-wise FDR-corrected *q* < 0.05, cluster size > 40). Regions outside the left putamen included the left middle occipital gyrus, right superior temporal gyrus, right supplementary motor area, right putamen, left precuneus, and right precentral gyrus. *B*, All clusters outside the left putamen showed no significant group difference in goal-dependency β estimates (*p*s > 0.18). Each dot represents an individual deaf signer (red = native; blue = delayed). MOG_L = left middle occipital gyrus; STG_R = right superior temporal gyrus; SMA_R = right supplementary motor area; PUT_R = right putamen; PCUN_L = left precuneus; PreCG_R = right precentral gyrus.

**Figure S7.**
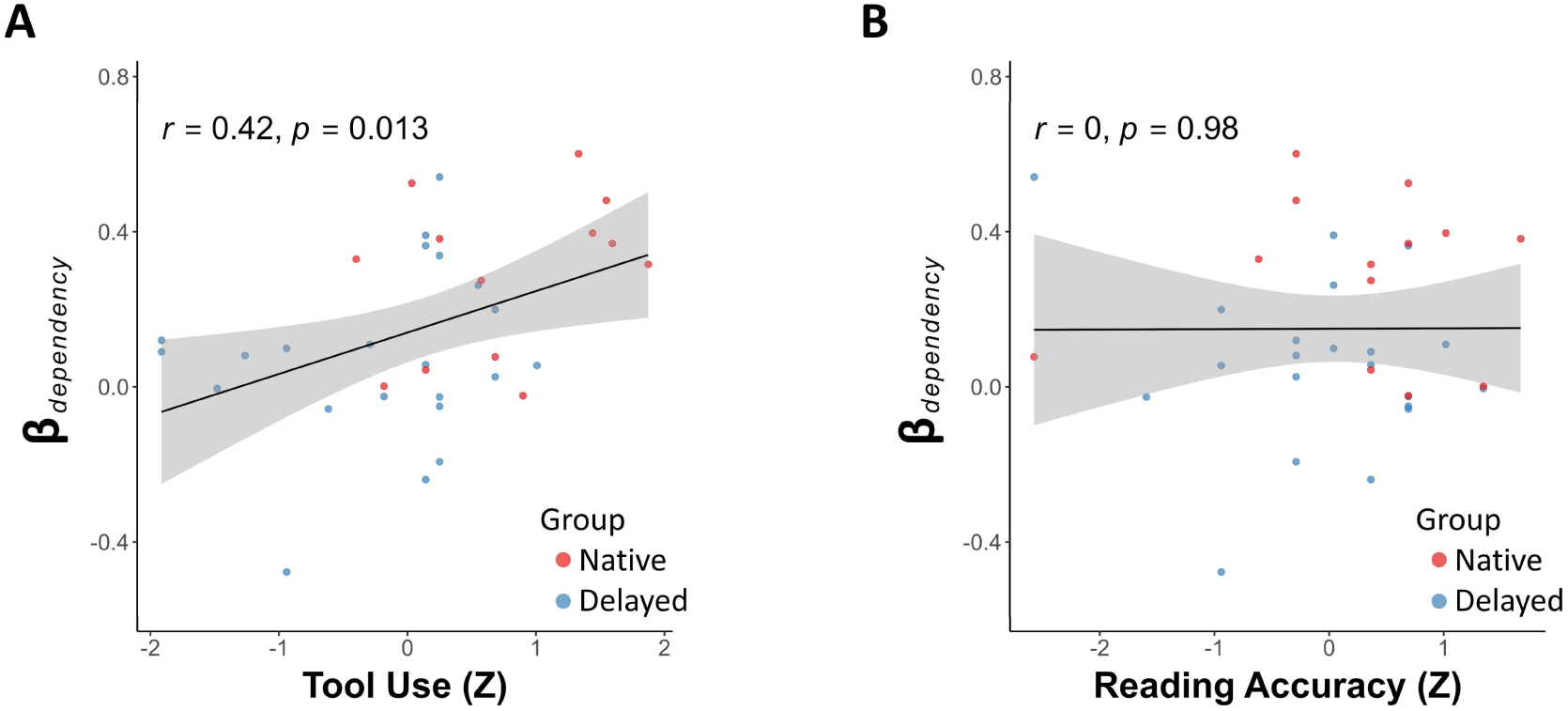
Putaminal encoding of goal dependency relates to tool use but not to Chinese reading. ***A***, Putaminal β values of goal dependency positively correlated with tool-use performance (*r* = 0.42, *p* = 0.013). Each dot represents an individual deaf signer (red = native; blue = delayed). ***B***, Putaminal β values of goal dependency were not correlated with Chinese reading accuracy (*r* = 0, *p* = 0.98). Each dot represents an individual deaf signer (red = native; blue = delayed). The regression line shows the fitted relationship, with shaded areas denoting the 95% confidence interval.

**Table S1.**
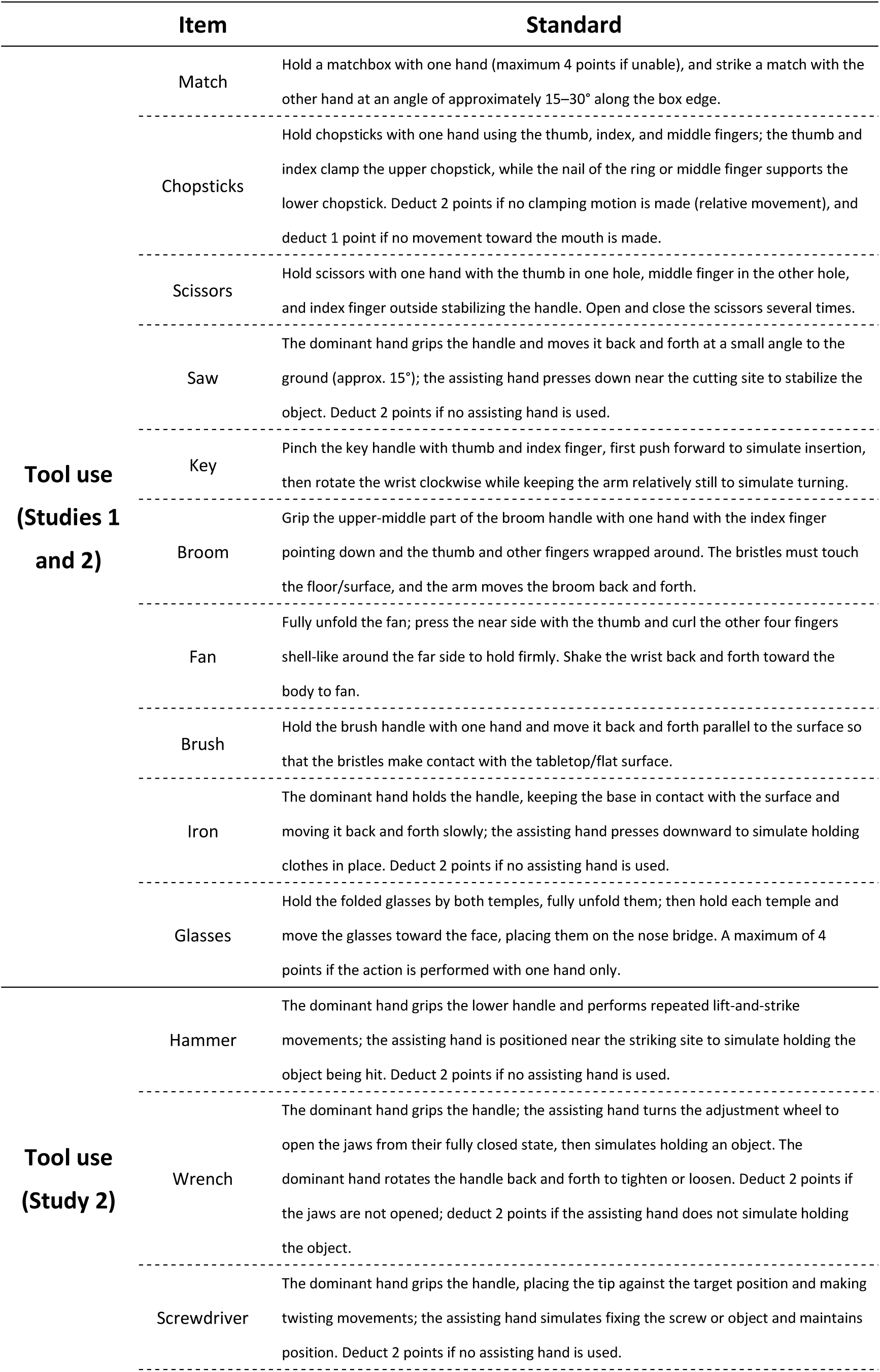
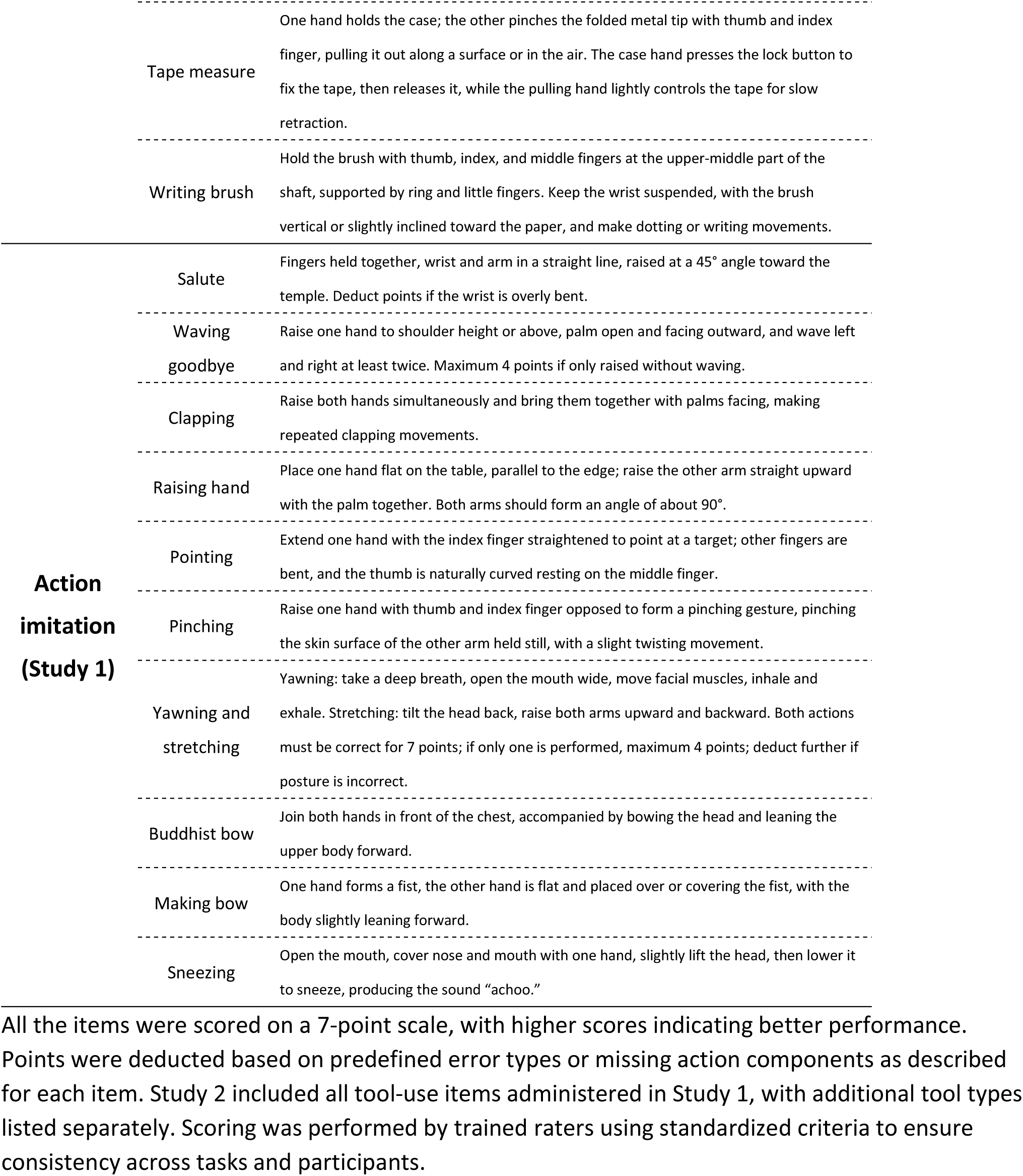
Scoring criteria for tool use and action imitation tasks.

**Table S2.** Collinearity diagnostics for the parametric modulators in Study 2.

(A) Pairwise correlations among modulators
| Pair | Pearson's r |
| --- | --- |
| Goal dependency – Element number | 0.64 |
| Goal dependency – Pixel number | –0.20 |
| Element number – Pixel number | 0.22 |

(B) Variance inflation factors (VIFs)
| Variable | VIF |
| --- | --- |
| Goal dependency | 2.11 |
| Element number | 2.12 |
| Pixel number | 1.32 |

Pairwise correlations and variance inflation factors (VIFs) were computed across object-level parametric modulator values, with one value per object entered into the fMRI general linear model (see Methods). All the VIF values were well below the commonly used thresholds for problematic multicollinearity.

## References

Alexander, G. E., DeLong, M. R., & Strick, P. L. (1986). Parallel Organization of Functionally Segregated Circuits Linking Basal Ganglia and Cortex. Annual Review of Neuroscience, 9(1), 357–381.

Arbib, M. A. (2011). From Mirror Neurons to Complex Imitation in the Evolution of Language and Tool Use. Annual Review of Anthropology, 40(1), 257–273. 10.1146/annurev-anthro-081309-145722

Arbuckle, J. L. (2011). IBM SPSS Amos 20 user’s guide. IBM SPSS.

Ash, S., Evans, E., O’Shea, J., Powers, J., Boller, A., Weinberg, D., Haley, J., McMillan, C., Irwin, D. J., Rascovsky, K., & Grossman, M. (2013). Differentiating primary progressive aphasias in a brief sample of connected speech. Neurology, 81(4), 329–336. 10.1212/WNL.0b013e31829c5d0e

Badre, D., & Frank, M. J. (2012). Mechanisms of Hierarchical Reinforcement Learning in Cortico-Striatal Circuits 2: Evidence from fMRI. Cerebral Cortex, 22(3), 527–536. 10.1093/cercor/bhr117

Bamford, N. S., Wightman, R. M., & Sulzer, D. (2018). Dopamine’s Effects on Corticostriatal Synapses during Reward-Based Behaviors. Neuron, 97(3), 494–510. 10.1016/j.neuron.2018.01.006

Bates, E., Wilson, S. M., Saygin, A. P., Dick, F., Sereno, M. I., Knight, R. T., & Dronkers, N. F. (2003). Voxel-based lesion–symptom mapping. Nature Neuroscience, 6(5), 448–450.

Bi, Y., Han, Z., Zhong, S., Ma, Y., Gong, G., Huang, R., Song, L., Fang, Y., He, Y., & Caramazza, A. (2015). The White Matter Structural Network Underlying Human Tool Use and Tool Understanding. The Journal of Neuroscience, 35(17), 6822–6835. 10.1523/JNEUROSCI.3709-14.2015

Bocanegra, Y., García, A. M., Pineda, D., Buriticá, O., Villegas, A., Lopera, F., Gómez, D., Gómez-Arias, C., Cardona, J. F., Trujillo, N., & Ibáñez, A. (2015). Syntax, action verbs, action semantics, and object semantics in Parkinson’s disease: Dissociability, progression, and executive influences. Cortex, 69, 237–254. 10.1016/j.cortex.2015.05.022

Bock, K., & Levelt, W. J. M. (Eds.). (1994). Language production: Grammatical encoding. In Handbook of Psycholinguistics (pp. 945–984). Academic Press.

Brozzoli, C., Roy, A., Lidborg, L. H., & Lövdén, M. (2019). Language as a Tool: Motor Proficiency Using a Tool Predicts Individual Linguistic Abilities. Frontiers in Psychology, 10. 10.3389/fpsyg.2019.01639

Buxbaum, L. J., Sirigu, A., Schwartz, M. F., & Klatzky, R. (2003). Cognitive representations of hand posture in ideomotor apraxia. Neuropsychologia, 41(8), 1091–1113. 10.1016/S0028-3932(02)00314-7

Caspers, S., Zilles, K., Laird, A. R., & Eickhoff, S. B. (2010). ALE meta-analysis of action observation and imitation in the human brain. NeuroImage, 50(3), 1148–1167. 10.1016/j.neuroimage.2009.12.112

Cataldo, D. M., Migliano, A., & Vinicius, L. (2018). Speech, stone tool-making and the evolution of language. PLoS ONE, 13. 10.1371/journal.pone.0191071

Chao, L. L., & Martin, A. (2000). Representation of Manipulable Man-Made Objects in the Dorsal Stream. NeuroImage, 12(4), 478–484. 10.1006/nimg.2000.0635

Chatham, C. H., & Badre, D. (2015). Multiple gates on working memory. Current Opinion in Behavioral Sciences, 1, 23–31. 10.1016/j.cobeha.2014.08.001

Choi, S., Na, Duk, Kang, E., Lee, K., Lee, S., & Na, Dong. (2001). Functional magnetic resonance imaging during pantomiming tool-use gestures. Experimental Brain Research, 139(3), 311–317. 10.1007/s002210100777

Collins, A. G. E., & Frank, M. J. (2013). Cognitive control over learning: Creating, clustering, and generalizing task-set structure. Psychological Review, 120(1), 190–229. 10.1037/a0030852

Coopmans, C. W., Kaushik, K., & Martin, A. E. (2023). Hierarchical structure in language and action: A formal comparison. Psychological Review, 130(4), 935–952. 10.1037/rev0000429

Copland, D. A., Brownsett, S., Iyer, K., & Angwin, A. J. (2021). Corticostriatal Regulation of Language Functions. Neuropsychology Review, 31(3), 472–494. 10.1007/s11065-021-09481-9

Fan, S., Wang, Xiaosha, Wang, Xiaoying, Wei, T., & Bi, Y. (2021). Topography of Visual Features in the Human Ventral Visual Pathway. Neuroscience Bulletin, 37(10), 1454–1468. 10.1007/s12264-021-00734-4

Fang, Y., Wang, X., Zhong, S., Song, L., Han, Z., Gong, G., & Bi, Y. (2018). Semantic representation in the white matter pathway. PLOS Biology, 16(4), e2003993. 10.1371/journal.pbio.2003993

Fazio, P., Cantagallo, A., Craighero, L., D’Ausilio, A., Roy, A. C., Pozzo, T., Calzolari, F., Granieri, E., & Fadiga, L. (2009). Encoding of human action in Broca’s area. Brain, 132(7), 1980–1988. 10.1093/brain/awp118

Folstein, M. F., Folstein, S. E., & McHugh, P. R. (1975). “Mini-mental state.” Journal of Psychiatric Research, 12(3), 189–198. 10.1016/0022-3956(75)90026-6

Frank, M. J., & Badre, D. (2012). Mechanisms of Hierarchical Reinforcement Learning in Corticostriatal Circuits 1: Computational Analysis. Cerebral Cortex, 22(3), 509–526. 10.1093/cercor/bhr114

Frank, M. J., Loughry, B., & O’Reilly, R. C. (2001). Interactions between frontal cortex and basal ganglia in working memory: A computational model. Cognitive, Affective, & Behavioral Neuroscience, 1(2), 137–160. 10.3758/CABN.1.2.137

Friederici, A. D. (2003). The Role of Left Inferior Frontal and Superior Temporal Cortex in Sentence Comprehension: Localizing Syntactic and Semantic Processes. Cerebral Cortex, 13(2), 170–177. 10.1093/cercor/13.2.170

Friederici, A. D., & Kotz, S. A. (2003). The brain basis of syntactic processes: Functional imaging and lesion studies. NeuroImage, 20, S8–S17. 10.1016/j.neuroimage.2003.09.003

Garcea, F. E., & Buxbaum, L. J. (2019). Gesturing tool use and tool transport actions modulates inferior parietal functional connectivity with the dorsal and ventral object processing pathways. Human Brain Mapping, 40(10), 2867–2883. 10.1002/hbm.24565

Gibson, E. (1998). Linguistic complexity: Locality of syntactic dependencies. Cognition, 68(1), 1–76. 10.1016/S0010-0277(98)00034-1

Giglio, L., Ostarek, M., Sharoh, D., & Hagoort, P. (2024). Diverging neural dynamics for syntactic structure building in naturalistic speaking and listening. Proceedings of the National Academy of Sciences, 121(11), e2310766121. 10.1073/pnas.2310766121

Goldenberg, G., & Randerath, J. (2015). Shared neural substrates of apraxia and aphasia. Neuropsychologia, 75, 40–49. 10.1016/j.neuropsychologia.2015.05.017

Goldenberg, G., & Spatt, J. (2009). The neural basis of tool use. Brain, 132(6), 1645–1655. 10.1093/brain/awp080

Graybiel, A. M. (1998). The Basal Ganglia and Chunking of Action Repertoires. Neurobiology of Learning and Memory, 70(1–2), 119–136. 10.1006/nlme.1998.3843

Greenfield, P. M. (1991). Language, tools and brain: The ontogeny and phylogeny of hierarchically organized sequential behavior. Behavioral and Brain Sciences, 14(4), 531–551. 10.1017/S0140525X00071235

Greenfield, P. M., & Westerman, M. A. (1978). Some psychological relations between action and language structure. Journal of Psycholinguistic Research, 7(6), 453–475. 10.1007/BF01068098

Haber, S. N. (2016). Corticostriatal circuitry. Dialogues in Clinical Neuroscience, 18(1), 7–21.

Han, Z., Ma, Y., Gong, G., He, Y., Caramazza, A., & Bi, Y. (2013). White matter structural connectivity underlying semantic processing: Evidence from brain damaged patients. Brain, 136(10), 2952–2965. 10.1093/brain/awt205

Harris, J. M., Saxon, J. A., Jones, M., Snowden, J. S., & Thompson, J. C. (2019). Neuropsychological differentiation of progressive aphasic disorders. Journal of Neuropsychology, 13(2), 214–239. 10.1111/jnp.12149

Hauser, M. D., Chomsky, N., & Fitch, W. T. (2002). The Faculty of Language: What Is It, Who Has It, and How Did It Evolve? Science, 298(5598), 1569–1579. 10.1126/science.298.5598.1569

Hazy, T. E., Frank, M. J., & O’Reilly, R. C. (2007). Towards an executive without a homunculus: Computational models of the prefrontal cortex/basal ganglia system. Philosophical Transactions of the Royal Society B: Biological Sciences, 362(1485), 1601–1613. 10.1098/rstb.2007.2055

Higuchi, S., Chaminade, T., Imamizu, H., & Kawato, M. (2009). Shared neural correlates for language and tool use in Broca’s area. NeuroReport, 20(15), 1376–1381. 10.1097/WNR.0b013e3283315570

Huang B., & Liao X. (Eds.). (2017). Modern Chinese (6th edition). Higher Education Press.

Iriki, A., & Taoka, M. (2012). Triadic (ecological, neural, cognitive) niche construction: A scenario of human brain evolution extrapolating tool use and language from the control of reaching actions. Philosophical Transactions of the Royal Society B: Biological Sciences, 367(1585), 10–23. 10.1098/rstb.2011.0190

Jackendoff, R. (2003). Précis of foundations of language: Brain, meaning, grammar, evolution. Behavioral and Brain Sciences, 26(6), 651–665. 10.1017/S0140525X03000153

Jin, X., & Costa, R. M. (2010). Start/stop signals emerge in nigrostriatal circuits during sequence learning. Nature, 466(7305), 457–462. 10.1038/nature09263

Jin, X., & Costa, R. M. (2015). Shaping action sequences in basal ganglia circuits. Current Opinion in Neurobiology, 33, 188–196. 10.1016/j.conb.2015.06.011

Jin, X., Tecuapetla, F., & Costa, R. M. (2014). Basal ganglia subcircuits distinctively encode the parsing and concatenation of action sequences. Nature Neuroscience, 17(3), 423–430. 10.1038/nn.3632

Johnson-Frey, S. H. (2004). The neural bases of complex tool use in humans. Trends in Cognitive Sciences, 8(2), 71–78. 10.1016/j.tics.2003.12.002

Kaas, J., & Stepniewska, I. (2023). The basal ganglia are a target for sensorimotor domains in posterior parietal, premotor, and motor cortex in primates. Current Opinion in Neurobiology, 83, 102783. 10.1016/j.conb.2023.102783

Kaufman, A. S. (2004). Kaufman Brief Intelligence Test–Second Edition (KBIT-2). American Guidance Service.

Klein, D., & Manning, C. D. (2003). Accurate unlexicalized parsing. Proceedings of the 41st Annual Meeting on Association for Computational Linguistics - ACL’03, 1, 423–430. 10.3115/1075096.1075150

Lewis, J. W. (2006). Cortical Networks Related to Human Use of Tools. The Neuroscientist, 12(3), 211–231. 10.1177/1073858406288327

Lombao, D., Guardiola, M., & Mosquera, M. (2017). Teaching to make stone tools: New experimental evidence supporting a technological hypothesis for the origins of language. Scientific Reports, 7(1), 14394. 10.1038/s41598-017-14322-y

Martin, A. (2007). The Representation of Object Concepts in the Brain. Annual Review of Psychology, 58(1), 25–45. 10.1146/annurev.psych.57.102904.190143

Martins, M. J. D., Bianco, R., Sammler, D., & Villringer, A. (2019). Recursion in action: An fMRI study on the generation of new hierarchical levels in motor sequences. Human Brain Mapping, 40(9), 2623–2638. 10.1002/hbm.24549

Mayberry, R. I., Chen, J.-K., Witcher, P., & Klein, D. (2011). Age of acquisition effects on the functional organization of language in the adult brain. Brain and Language, 119(1), 16–29. 10.1016/j.bandl.2011.05.007

Mayberry, R. I., Lock, E., & Kazmi, H. (2002). Linguistic ability and early language exposure. Nature, 417(6884), 38–38. 10.1038/417038a

Miller, L. E., Montroni, L., Koun, E., Salemme, R., Hayward, V., & Farnè, A. (2018). Sensing with tools extends somatosensory processing beyond the body. Nature, 561(7722), 239–242. 10.1038/s41586-018-0460-0

Morgan, T. J. H., Uomini, N. T., Rendell, L. E., Chouinard-Thuly, L., Street, S. E., Lewis, H. M., Cross, C. P., Evans, C., Kearney, R., de la Torre, I., Whiten, A., & Laland, K. N. (2015). Experimental evidence for the co-evolution of hominin tool-making teaching and language. Nature Communications, 6(1), 6029. 10.1038/ncomms7029

Moro, A. (2014). On the similarity between syntax and actions. Trends in Cognitive Sciences, 18(3), 109–110. 10.1016/j.tics.2013.11.006

Moro, A., Tettamanti, M., Perani, D., Donati, C., Cappa, S. F., & Fazio, F. (2001). Syntax and the Brain: Disentangling Grammar by Selective Anomalies. NeuroImage, 13(1), 110–118. 10.1006/nimg.2000.0668

Napoli, D. J., & Sutton-Spence, R. (2014). Order of the major constituents in sign languages: Implications for all language. Frontiers in Psychology, 5. 10.3389/fpsyg.2014.00376

Newport, E. L. (1990). Maturational Constraints on Language Learning. Cognitive Science, 14(1), 11–28. 10.1207/s15516709cog1401_2

Nicola, S. M., Surmeier, D. J., & Malenka, R. C. (2000). Dopaminergic Modulation of Neuronal Excitability in the Striatum and Nucleus Accumbens. Annual Review of Neuroscience, 23(1), 185–215. 10.1146/annurev.neuro.23.1.185

O’Reilly, R. C., & Frank, M. J. (2006). Making Working Memory Work: A Computational Model of Learning in the Prefrontal Cortex and Basal Ganglia. Neural Computation, 18(2), 283–328. 10.1162/089976606775093909

Pastra, K., & Aloimonos, Y. (2012). The minimalist grammar of action. Philosophical Transactions of the Royal Society B: Biological Sciences, 367(1585), 103–117. 10.1098/rstb.2011.0123

Peirce, J., Gray, J. R., Simpson, S., MacAskill, M., Höchenberger, R., Sogo, H., Kastman, E., & Lindeløv, J. K. (2019). PsychoPy2: Experiments in behavior made easy. Behavior Research Methods, 51(1), 195–203. 10.3758/s13428-018-01193-y

Pulvermüller, F. (2014). The syntax of action. Trends in Cognitive Sciences, 18(5), 219–220. 10.1016/j.tics.2014.01.001

Py, R., Grosbras, M.-H., Brozzoli, C., & Montant, M. (2025). A tool to probe domain-general syntax: Simple and complex actions with a tool improve syntactic comprehension in language. Current Research in Behavioral Sciences, 9, 100190. 10.1016/j.crbeha.2025.100190

Rizzolatti, G., & Arbib, M. A. (1998). Language within Our Grasp. Trends in Neurosciences, 21(5), 188–194. 10.7551/mitpress/3077.003.0020

Rochon, E., Saffran, E. M., Berndt, R. S., & Schwartz, M. F. (2000). Quantitative Analysis of Aphasic Sentence Production: Further Development and New Data. Brain and Language, 72(3), 193–218. 10.1006/brln.1999.2285

Rosenbaum, D. A., Marchak, F., Barnes, H. J., Vaughan, J., Slotta, J. D., & Jorgensen, M. J. (1990). Constraints for Action Selection: Overhand Versus Underhand Grips. In M. Jeannerod (Ed.), Attention and Performance XIII (pp. 359–386). Psychology Press.

Rosenbaum, D. A., Meulenbroek, R. J., Vaughan, J., & Jansen, C. (2001). Posture-based motion planning: Applications to grasping. Psychological Review, 108(4), 709–734. 10.1037/0033-295X.108.4.709

Roy, A. C., & Arbib, M. A. (2005). The syntactic motor system. Gesture, 5(1–2), 7–37. 10.1075/gest.5.1.03roy

Sandler, W., & Lillo-Martin, D. C. (2006). Sign language and linguistic universals. Cambridge university press.

Seger, C. A. (2008). How do the basal ganglia contribute to categorization? Their roles in generalization, response selection, and learning via feedback. Neuroscience & Biobehavioral Reviews, 32(2), 265–278. 10.1016/j.neubiorev.2007.07.010

Soni, A., & Frank, M. J. (2025). Adaptive chunking improves effective working memory capacity in a prefrontal cortex and basal ganglia circuit. eLife, 13, RP97894. 10.7554/eLife.97894

Steele, J., Ferrari, P. F., & Fogassi, L. (2012). From action to language: Comparative perspectives on primate tool use, gesture and the evolution of human language. Philosophical Transactions of the Royal Society B: Biological Sciences, 367(1585), 4–9. 10.1098/rstb.2011.0295

Stout, D., & Chaminade, T. (2012). Stone tools, language and the brain in human evolution. Philosophical Transactions of the Royal Society B: Biological Sciences, 367(1585), 75–87. 10.1098/rstb.2011.0099

Tan Y., Ye Z., Zhang R., & Zhou X. (2026). The Role of Basal Ganglia in Language Comprehension. Journal of Psychological Science, 49(1), 238–251. 10.16719/j.cnki.1671-6981.20260121

Thibault, S., Py, R., Gervasi, A. M., Salemme, R., Koun, E., Lövden, M., Boulenger, V., Roy, A. C., & Brozzoli, C. (2021). Tool use and language share syntactic processes and neural patterns in the basal ganglia. Science, 374(6569), eabe0874. 10.1126/science.abe0874

Tzourio-Mazoyer, N., Landeau, B., Papathanassiou, D., Crivello, F., Etard, O., Delcroix, N., Mazoyer, B., & Joliot, M. (2002). Automated Anatomical Labeling of Activations in SPM Using a Macroscopic Anatomical Parcellation of the MNI MRI Single-Subject Brain. NeuroImage, 15(1), 273–289. 10.1006/nimg.2001.0978

Ullman, M. T. (2001). A neurocognitive perspective on language: The declarative/procedural model. Nature Reviews Neuroscience, 2(10), 717–726. 10.1038/35094573

Ullman, M. T. (2004). Contributions of memory circuits to language: The declarative/procedural model. Cognition, 92(1–2), 231–270. 10.1016/j.cognition.2003.10.008

Vaesen, K. (2012). The cognitive bases of human tool use. Behavioral and Brain Sciences, 35(4), 203–218. 10.1017/S0140525X11001452

Vingerhoets, G., Alderweireldt, A.-S., Vandemaele, P., Cai, Q., Van der Haegen, L., Brysbaert, M., & Achten, E. (2013). Praxis and language are linked: Evidence from co-lateralization in individuals with atypical language dominance. Cortex, 49(1), 172–183. 10.1016/j.cortex.2011.11.003

Wang, X., Wang, B., & Bi, Y. (2023). Early language exposure affects neural mechanisms of semantic representations. eLife, 12, e81681. 10.7554/eLife.81681

Weiss, P. H., Ubben, S. D., Kaesberg, S., Kalbe, E., Kessler, J., Liebig, T., & Fink, G. R. (2016). Where language meets meaningful action: A combined behavior and lesion analysis of aphasia and apraxia. Brain Structure and Function, 221(1), 563–576. 10.1007/s00429-014-0925-3

Wen, H., Wang, D., & Bi, Y. (2024). Processing Language Partly Shares Neural Genetic Basis with Processing Tools and Body Parts. Eneuro, 11(8), ENEURO.0138-24.2024. 10.1523/ENEURO.0138-24.2024

Yan, C.-G., Wang, X.-D., Zuo, X.-N., & Zang, Y.-F. (2016). DPABI: Data Processing & Analysis for (Resting-State) Brain Imaging. Neuroinformatics, 14(3), 339–351. 10.1007/s12021-016-9299-4

Yang, Z., Inagaki, M., Gerfen, C. R., Fontolan, L., & Inagaki, H. K. (2025). Integrator dynamics in the cortico-basal ganglia loop for flexible motor timing. Nature, 1–10. 10.1038/s41586-025-09778-2

